# Molecular Pixelation: Single cell spatial proteomics by sequencing

**DOI:** 10.1101/2023.06.05.543770

**Authors:** Filip Karlsson, Tomasz Kallas, Divya Thiagarajan, Max Karlsson, Maud Schweitzer, Jose Fernandez Navarro, Louise Leijonancker, Sylvain Geny, Erik Pettersson, Jan Rhomberg-Kauert, Marcela Gonzalez Granillo, Jessica Bunz, Johan Dahlberg, Michele Simonetti, Prajakta Sathe, Petter Brodin, Alvaro Martinez Barrio, Simon Fredriksson

## Abstract

The spatial distribution of cell surface proteins govern vital processes of the immune system such as inter-cell communication and mobility. However, tools for studying these at high multiplexing scale, resolution, and throughput needed to drive novel discoveries are lacking. We present Molecular Pixelation, a DNA-sequencing based method for single cell analysis to quantify protein abundance, spatial distribution, and colocalization of targeted proteins using Antibody Oligonucleotide Conjugates (AOCs). Relative locations of AOCs are inferred by sequentially associating these into local neighborhoods using DNA-pixels containing unique pixel identifier (UPI) sequences, forming >1,000 connected spatial zones per single cell in three dimensions. DNA-sequencing reads are computationally arranged into spatial single cell maps for 76 proteins without cell compartmentalization. By studying immune cell dynamics and using spatial statistics on graph representations of the data, previously known and novel patterns of protein spatial polarization and co-localization were found in chemokine-stimulated T-cells.

## Introduction

The spatial organization of immune cell surface receptors governs multiple functions such as dynamical tuning of cell signaling (1), cell-cell communication (2), T-cell effector function (3), movement via adhesion receptors (4), drug mode-of-action (5), and efficacy of cellular therapies (6). Spatial protein organization has traditionally been studied with antibodies carrying fluorophores using fluorescence microscopy or Imaging Flow Cytometry (7), typically providing data in one focal plane. The generation of three dimensional data at high throughput is thus limited by the need for microscopy imaging of one focal plane at a time for each fluorophore/antibody combination. Signal to noise is also hampered by auto-flourescence and spectral bleed-through between channels. Super-resolution imaging methods have provided groundbreaking insights into cellular functional responses and signaling (8), but are limited in multiplexing and throughput. Single cell targeted proteomics with multiplexing levels higher than achievable with fluorophores can be enabled by tagging antibodies with DNA (9) but requires single cell compartmentalization and does not provide any spatial information. The promise of using methods solely relying on nucleic acid sequence to image biological samples has been highlighted (10,11) and demonstrated for RNA in 2D imaging (12). These methods, at times referred to as DNA-Microscopy, where multiple DNA-tag types reflect biomolecule identity as well as a position in the biological sample, could increase sample throughput, multiplexing and potentially resolution.

Here we present Molecular Pixelation (MPX) that uses DNA-tagged antibodies (Antibody-Oligonucleotide Conjugates, AOCs) bound to their protein targets on chemically fixed cells to survey cell surface protein arrangement and dynamics in a highly multiplex fashion. The assay is performed in solution in a single reaction tube, without the need to compartmentalize each cell. Spatial analysis of protein arrangement is enabled by serially forming associations between spatially proximate AOCs into local neighborhoods via the incorporation of a unique molecular identifier (UMI), similar to the Proximity Barcoding Assay (13). The generated amplicons are sequenced and spatial relationships of proteins are inferred from graph representations of the data for each single cell.

The association of AOCs bound to cells into local neighborhoods is done using so-called DNA-Pixels, which are single-stranded DNA molecules, each containing a concatemer of a unique molecular identifier sequence called a Unique Pixel Identifier (UPI) generated by rolling circle amplification from circular DNA templates. Once added to the reaction, each DNA pixel can hybridize to multiple AOC molecules in proximity on the cell surface. The UPI sequence of the hybridized DNA pixel is then incorporated onto the AOC oligonucleotide by a gap-fill ligation reaction, thus creating neighborhoods where the set of AOCs within each neighborhood share the same UPI sequence. Following enzymatic degradation of the first DNA pixel set, a second set of DNA pixels is similarly incorporated by hybridization and gap-fill ligation reactions (Figure 1). The generated amplicons are then amplified by PCR and sequenced. Each sequenced molecule contains four distinct DNA barcode motifs; a unique molecule identifier (UMI) to enable identification of unique AOC molecules, a protein identity barcode, and two UPI barcodes with neighborhood memberships.

**Figure 1.**
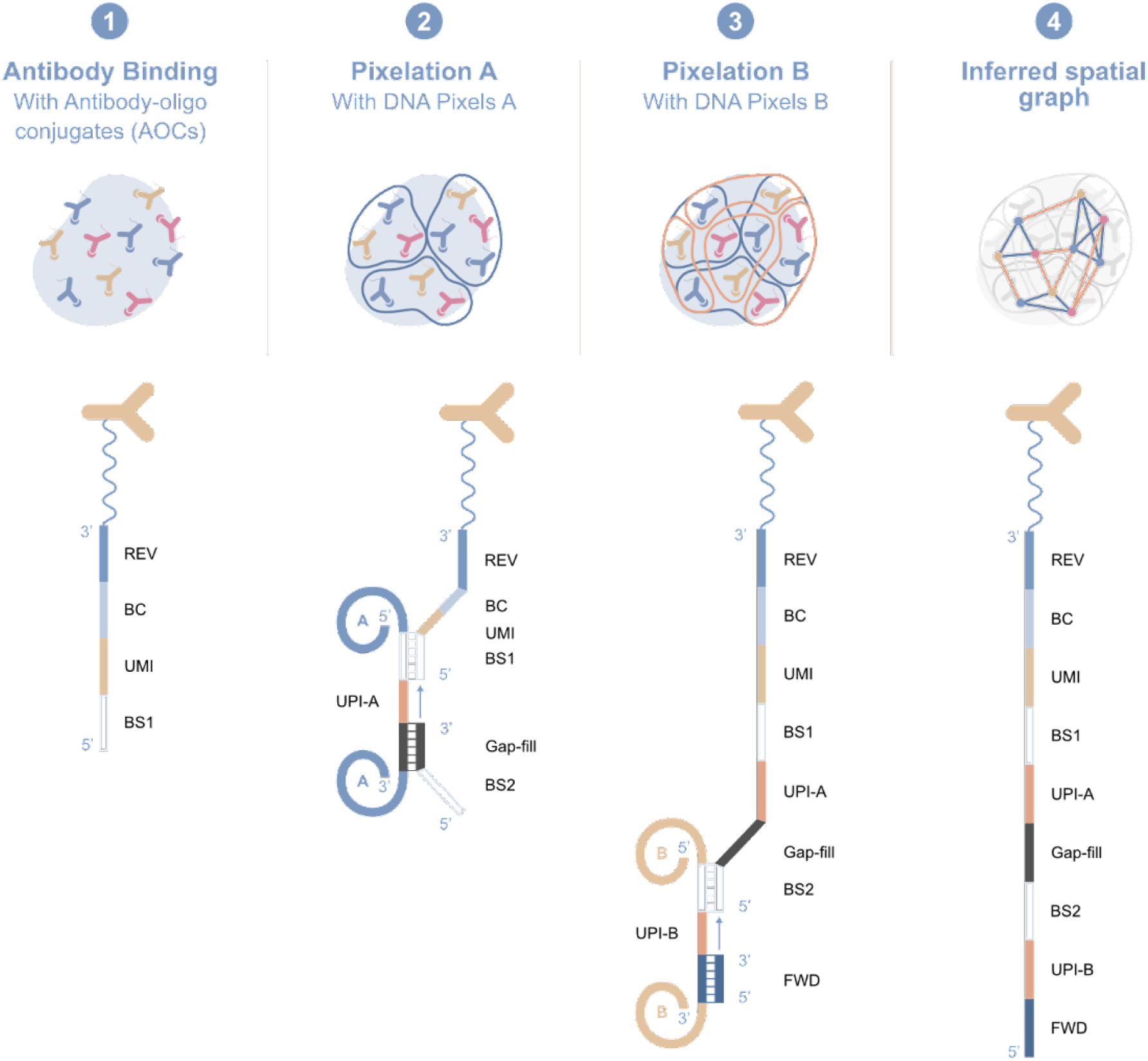
Molecular Pixelation. 1) AOCs, containing a protein identifying sequence (BC), a unique molecule identifier (UMI) sequence and a DNA pixel binding sequence (BS1) are bound to target proteins on a cell. 2) DNA pixel set A then hybridizes to AOCs, followed by gap-fill ligation to incorporate the unique pixel identifier (UPI) sequence of each DNA pixel and a common pixel binding site 2 (BS2) onto the 5’ end of each AOC, dividing each cell into >1,000 zones where all AOCs within each zone share the same UPI-A sequence. 3) Similarly, DNA-pixel set B hybridizes to the extended AOCs and the UPI-B sequence is incorporated using gap-fill ligation. The final product is amplified by PCR and sequenced. 4) Each sequenced AOC molecule is represented as an edge in a bipartite graph, with the two UPI sequences as nodes, and protein identities as either edge or node attributes. The sequencing data representing each cell is thereby organized into separated connected graph components from the association of AOC molecules into two partially overlapping UPI zones. Spatial analysis of protein arrangement is enabled by interrogating the location of node or edge attributes within the graph.

The relative location of each unique AOC molecule can be inferred from the overlap of UPI neighborhoods created from the two serial DNA pixel hybridization and gap-fill ligation steps (Figure 1). Each sequenced unique molecule can be represented as an edge in a bipartite graph, with UPI-A and UPI-B sequences as nodes and protein identity as edge attributes, or alternatively as a one-mode projected graph of UPI-A sequences as nodes and protein identities as node attributes. The graphs generated from a sequenced sample following data processing and filtering, contains graph components which can be separated into distinct graphs per single cell. Spatial analysis of protein arrangement, such as the degree of clustering of a single protein or colocalization between two or more proteins can be performed by interrogating the location of edge or node attributes on the graph representations of each cell.

Using a panel of 76 AOCs targeting immune cell surface proteins, and four control AOCs, we demonstrate the ability of MPX to generate single-cell data based on protein abundance from PBMCs. Next, the method was used to quantify the degree of spatial clustering or polarization from Polarity scores of each assayed protein upon modulation of the cell by a therapeutic antibody or by capping using a secondary antibody. Finally, abundance, polarity score and pairwise colocalization of the target proteins was studied on immune cells subjected to chemotactic migration stimulation.

## Results

MPX data comprises both the spatial location and the abundance of measured proteins, and can thus be processed similarly to data from other single cell technologies to annotate cells by their identity, as defined by the proteins displayed on their cell surface. To demonstrate the ability to generate single cell data with MPX, without the need to compartmentalize individual cells, a heterogeneous sample was processed using a 76-plex target panel against T cells, NK cells, B cells and monocytes and from the generated count data the distribution of protein count signatures generated from each cell was assessed.

### Cell annotation of PBMC from AOC count data

Peripheral blood mononuclear cells (PBMCs) from a healthy donor were fixed with Paraformaldehyde (PFA) and assayed in two replicates by first staining with AOCs, performing MPX, and taking a subset of 500 cells to PCR-based library preparation for sequencing. After data processing of sequence reads using the Pixelator pipeline (Methods), 468 and 553 distinct cells were identified in the output data for the two replicates, which correlated well with the 500 cells input to PCR and sequencing. Titration of cell input numbers between 50 to 1000 cells has further shown that there is a strong correlation between the number of cells input for PCR to the number of detected cells in data following Pixelator processing, indicating that one and not several connected graph components are generated from each cell following data processing. (Supplementary Figure S1).

The MPX protein count data matrices were processed to visualize and annotate the cellular identities across the cells in the two replicate samples (see Methods). Relative cell counts were scaled and log-normalized, and used to identify 6 clusters of cells with shared identity, using Louvain community detection on a Shared Nearest Neighbor graph. The cells were visualized in a Uniform Manifold Approximation and Projection (UMAP) (14) (Figure 2A), which displays separated clusters corresponding to the main cell populations expected in PBMCs; T cells, B cells, NK cells, and monocytes. This annotation of cells was established using a differential abundance analysis (Wilcoxon Rank Sum test, downsampled to 50 cells per cluster), where each cluster was compared to other cells, resulting in 3 to 31 proteins with significantly different abundance (Bonferroni adjusted p-value < 0.01, absolute average log2 Fold Change > 1) per cluster (Figure 2B). The formation of separate cell-type specific clusters demonstrates that MPX is able to perform single-cell profiling without the need for physical compartmentalization of each individual cell nor combinatorial cell barcoding using split-pool strategies. Although MPX data contained a low level of incompatible markers, this was estimated to be in range with observed rates from other single-cell methods (15).

**Figure 2.**
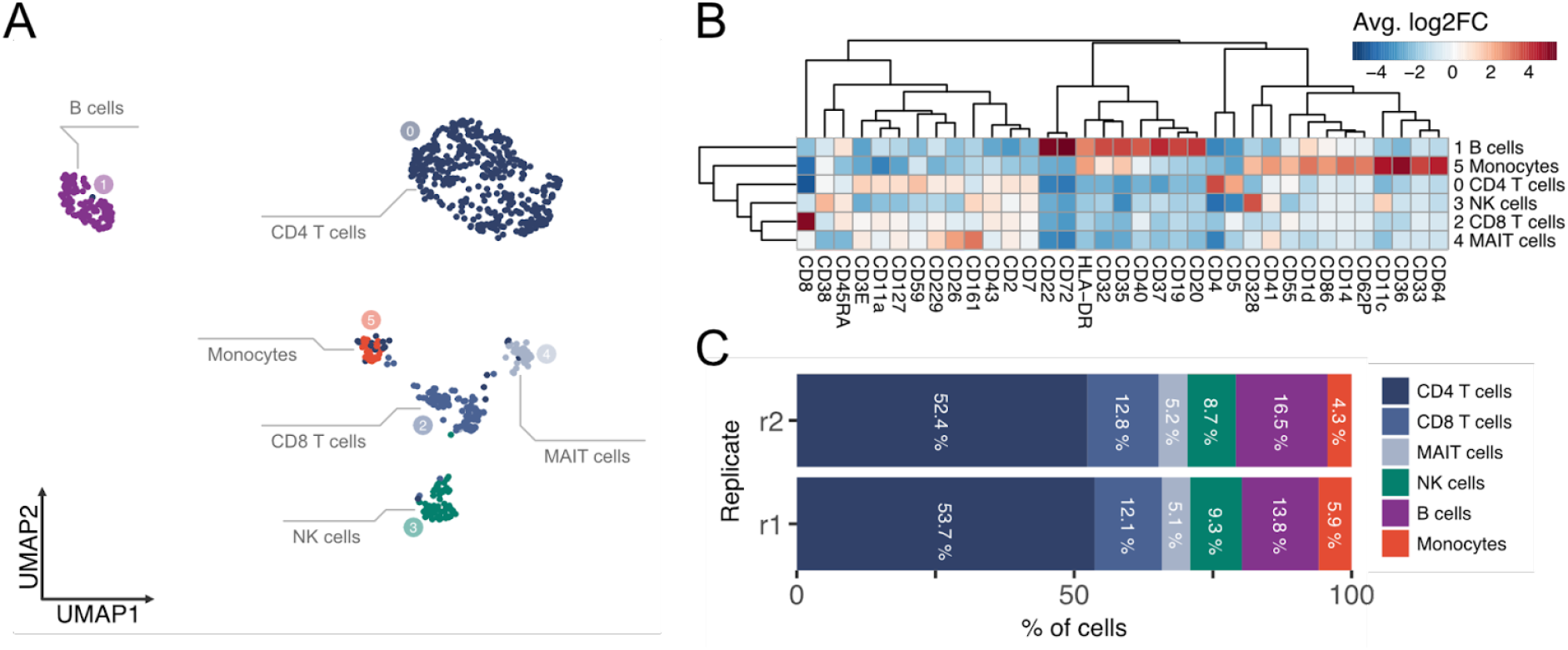
Count data from an MPX experiment of healthy PBMCs; A) UMAP of PBMC following Molecular Pixelation. The connected graph components representing each cell clustered into separated groups representing the major cell types in PBMCs. B) Heatmap with relative expression of differentially abundant proteins across the 6 clusters of cell identities, showing the average log2 fold change of proteins for the cluster compared to other cells. Proteins with significantly different abundance (Bonferroni adjusted p-value < 0.01, and absolute average log2 fold change > 2) for at least one cluster are shown. C) Frequencies of annotated cell types per replicate.

Abundance from all targets showed expected specificity patterns, exemplified by CD3 cooccurrent with CD4 or CD8 in CD4 T cells and CD8 T cells respectively, and by CD19 and CD20 exclusively abundant on B cells (Supplementary Figure S2). Isotype control levels were low, with mIgG1, mIgG2a, and mIgG2b constituting only 0.03%, 0.15% and 0.04% respectively of the total AOC counts per cell on average (Supplementary Figure S3). The cytoplasmic control target beta actin (ACTB), used to verify plasma membrane integrity, also showed low levels.

The estimated frequencies of the identified cell populations were similar between the two replicates samples, with average levels around expected percentages; 70.6% T cells (52.9% CD4 T, 12.5% CD8 T, 5.2% MAIT), 9.0% NK cells, 15.4% B cells, and 5.0% Monocytes (Figure 2C). MPX generated on average 1,742 DNA-pixel A zones and 9,600 AOC UMIs per cell, with 5.6 UMIs per UPI-A pixel (Supplementary Figure S4).

### Spatial autocorrelation from stimulated T and B cells

The graph-based data generated by MPX can be used for spatial analysis by interrogating the edge or node attributes representing different protein targets. Spatial autocorrelation is a measure that can be used to measure clustering or non-randomness of a spatial variable. A Polarity score, derived from the Moran’s I autocorrelation statistic can thus be calculated for each protein marker per cell from spatial weights derived from the adjacency matrix of cell graphs (see Methods), where positive Polarity scores indicate clustered spatial distribution and scores centered around zero indicate random spatial distribution.

To evaluate the ability to detect clustered protein expression from MPX data, we spatially clustered, or “polarized” CD3 by a capping reaction, using the CD3 AOC and a secondary anti-mouse antibody prior to PFA fixation, staining with the remaining AOCs, and MPX. Data generated from 3 separate experiments were combined for a total of 2075 and 1359 cells after data filtering for the capped and control samples respectively. In a second approach to demonstrate protein polarization, we analyzed the distribution of CD20 after treatment of Raji cells with a Rituximab-based AOC before PFA fixation. Rituximab, a monoclonal therapeutic antibody, is known to cluster CD20 on B-cells enhancing antibody dependent cellular cytotoxicity based cancer killing (5). Raji cells, not treated with Rituximab prior to fixation, were used as a negative control.

As seen in Figure 3A and 3D, Polarity scores were significantly elevated for CD3 capped and the Rituximab treated samples (Wilcoxon Rank Sum test downsampled to 50 cells; Benjamini Hochberg (BH) adjusted p-value of 2.0 × 10^−14^ and 1.2 × 10^−14^, respectively) compared to controls. No other proteins showed similar levels of difference between the treated sample and control, as can be seen in the resulting volcano plots (Supplementary Figures S5). Fluorescent microscopy for CD3 and Rituximab was performed to validate the spatial redistribution of the target proteins upon stimulation (Figure 3E, 3F). Spherical 3D density heat maps were generated from force-directed graph layouts of one representative CD3-capped T cell (Polarity Score 25) and one CD20/Rituximab-treated Raji cell (Polarity Score 54) to visualize the polarized distribution (Figure 3B & 3E).

**Figure 3.**
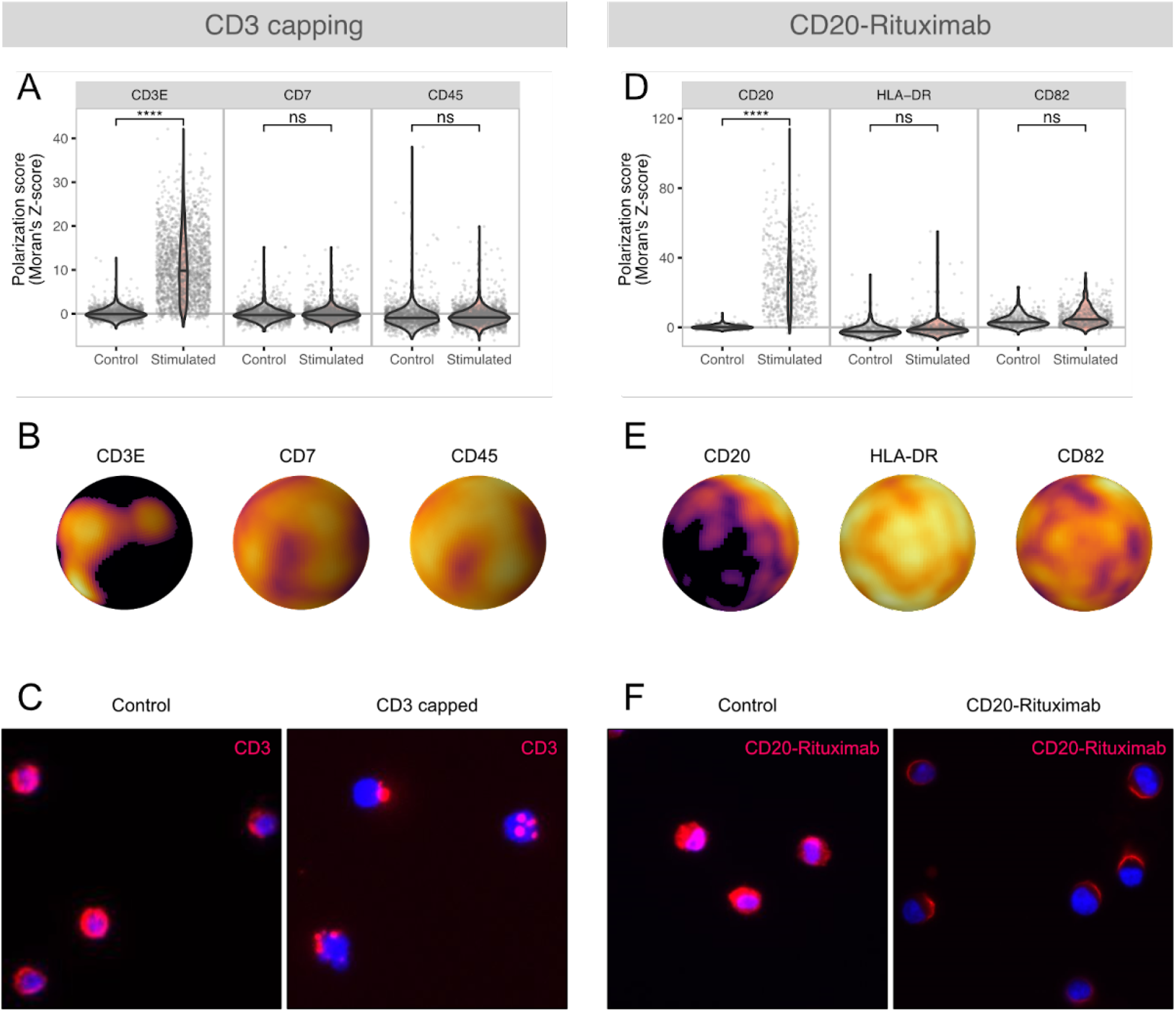
Protein polarization triggered by cell stimulation detected by spatial Polarity scores in MPX data for CD3-capped T-cells and Rituximab treated Raji B-cells. A,D) Violin plot of Polarity scores for CD3 and CD20/Rituximab, as well as for additional examples. B,E) 3D density heat maps derived from the graph representations of a CD3-capped cell (B) and a Rituximab treated cell (E), colored by the count density of the indicated markers. The higher density areas are of yellow color, while low density areas are dark. C,F) Fluorescence microscopy for CD3 and Rituximab in control and treated samples validated the presence of spatially clustered distribution for CD3 and Rituximab respectively in treated samples compared to controls.

For the Rituximab experiment, positive Polarity scores were also found for protein markers CD54 and CD82, which was observed in both negative controls and treated samples, indicating that these proteins have a clustered spatial distribution on Raji cells. Data for all detected targets are found in Supplementary Figure S6.

### Colocalization of protein pairs in chemotactic T-cells

Colocalization of cell surface proteins plays a crucial role in driving various cellular processes. To accurately quantify pairwise combinations of measured markers and identify patterns of colocalized protein groups, we developed a colocalization score (see Methods). This score is based on Pearson’s correlation coefficient (r) and permutation testing, and enables the comparison of measured scores with scores obtained from simulated cells that possess an equal distribution of marker counts but random localization. By doing so, the score reflects the degree of subcellular spatial co-occurrence of two proteins, representing the deviation from what would be expected by random chance. Essentially, it measures how much higher or lower a protein pair colocalizes compared to what would be predicted by chance, taking into account the frequency of each protein’s presence in the graph data. This approach minimizes the influence of experimental perturbations that may affect protein abundance and potentially inflate colocalization scores.

To demonstrate the applicability of our approach, we showcase the application of MPX in detecting cellular structures, specifically focusing on uropod formation in migrating immune cells. Uropods are critical for cytotoxic T-cells to infiltrate tumors (16), and their formation is associated with immune checkpoint inhibition efficacy and overall cancer survival (17). The high multiplexing capability of our method, coupled with the graph-based data it generates, facilitates unbiased discovery of colocalization patterns and even the segregation of protein pair localizations.

Two different types of conditions were applied to stimulate T cells to attain a migratory phenotype and form uropods. The cells were exposed to a plate immobilized with either CD54 (ICAM1) alone or in combination with a chemokine; either CCL2 (MCP1) or CCL5 (RANTES), triggering cell adhesion to the surface and uropod formation. The experiment consisted of six samples, assessing each combination of the two conditions: 1) CD54 immobilized surface or in solution, and 2) MCP1, RANTES, or unstimulated.

The effects of the conditions were analyzed from three distinct perspectives: protein abundance, polarization, and colocalization. The unstimulated non-coated sample served as the reference condition, representing minimal uropod formation. Statistical significance was assessed using the Wilcoxon Rank Sum test to determine if there were significant differences in protein abundance, polarization scores, and protein-protein colocalization scores between each condition and the control sample.

The analysis of protein abundance and spatial distribution revealed noteworthy results, with the most pronounced differences observed in the samples that were both CD54 immobilized and chemokine-treated, compared to the control. Interestingly, minimal effects were observed for the samples stimulated in solution, with limited changes in protein abundance, polarization, and colocalization under these conditions. The differential abundance analysis resulted in 12 proteins with significantly different protein abundance (BH adjusted p < 0.05, average log_2_ fold change > 0.5) in any of the conditions (Figure 4A). The differential polarization analysis identified 15 markers exhibiting significant differentiation in protein polarization (BH adjusted p < 0.05, average difference > 0.0125; Figure 4B), and the differential colocalization analysis identified 15 markers displaying significant differences in protein colocalization (BH adjusted p < 0.05, diff > 0.75; Figure 4C. Importantly, a notable overlap was observed between the markers demonstrating differential colocalization and those showing differential polarization, with 10 proteins displaying significant differences in both spatial tests across at least one of the conditions. Among these proteins, CD2, CD44, CD50, and CD162, which are known to be located within the T cell uropod (18), exhibited differential spatial arrangement in CD54 immobilized and stimulated samples, with increased polarization and increased colocalization between CD162 and CD50 (Figure 4D). Notably, CD37 has previously been described as participating in uropod formation in B cells, neutrophils, and dendritic cells (19), and is here observed to increase in polarization, and colocalization with CD50 and CD162 upon CD54 immobilization and chemokine stimulation in T-cells. Additionally, CD43, CD11a, and CD18, which are ligands of CD54 used in the stimulation/attachment of the T-cells (18), show decreased abundance (CD43 and CD11a) and increased polarization (CD11a and CD18), which may be effects of the CD54 immobilization partially blocking or perturbing AOC binding. In Figure 4E (and Supplementary Figures S7 & S8), representative cells from the sample with immobilized cells treated with RANTES displayed distinct spatial patterns of CD50, CD37, and CD162, characterized by colocalization and polarization, implicating the uropod. The polarization and pronounced three-way colocalization among CD50, CD37, and CD162 is visualized as a graph representation in Figure 4F (and Supplementary Figure S9).

**Figure 4.**
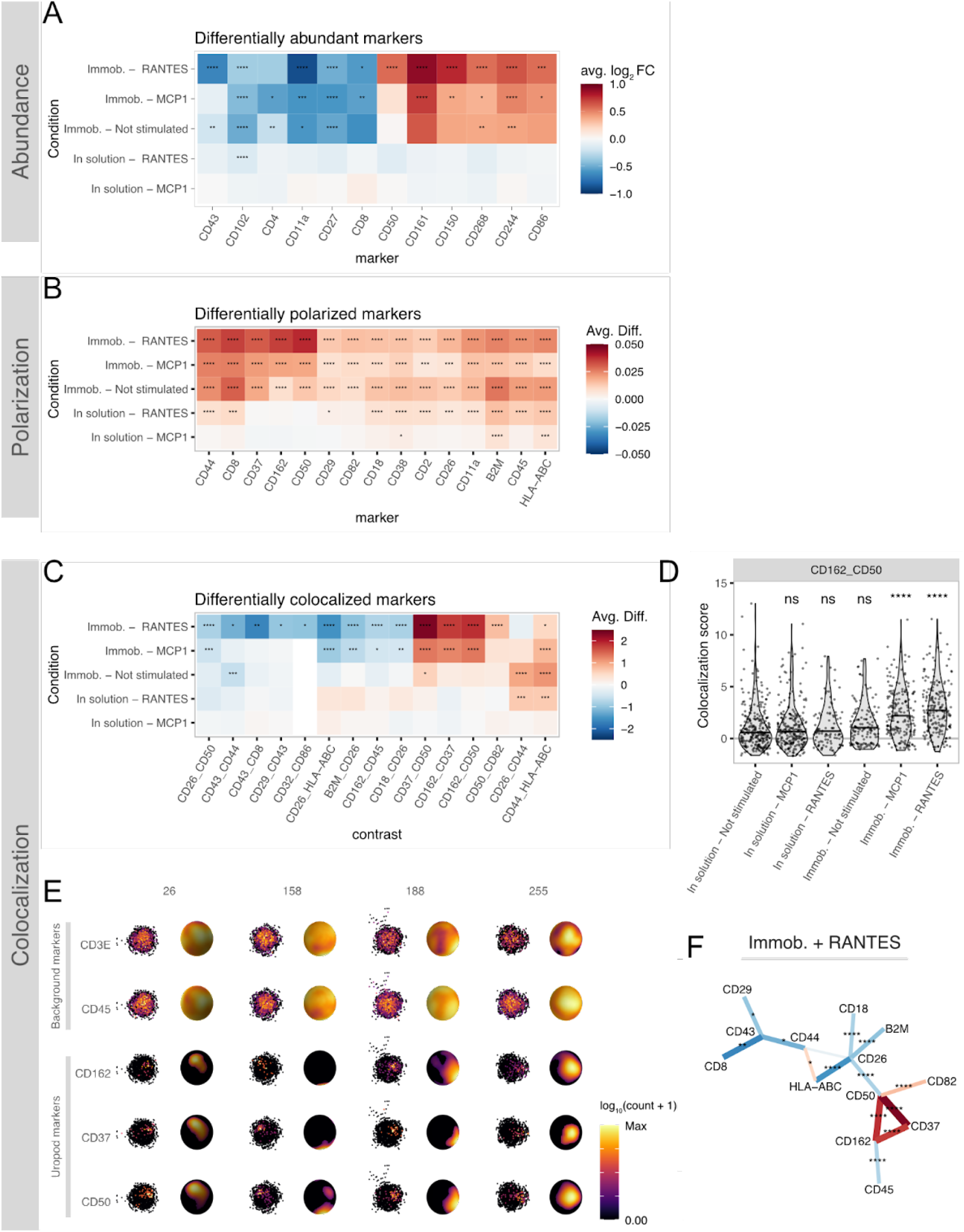
MPX analysis of chemokine stimulation of CD54 (ICAM1) immobilized T-cells. A-C) Heatmaps showing the effect sizes in differential analysis comparing abundance levels (A), Polarity scores (B), and colocalization scores (C) of each condition to the control sample of unstimulated T cells in suspension. In A, the color of the tile encodes the estimated log_2_ Fold change in abundance, while B and C show the difference in mean score. P-value levels are denoted: “*” = p < 0.05, “**” = p < 0.01, “***” = p < 0.001, “****” = p < 0.001. D) Pairwise colocalization score for uropod proteins CD162 and CD50 across all the sample conditions for each single cell. E) Visualization of 4 individual single cells (cell ID’s are denoted at the top of each column) showing 2 ubiquitous background markers (CD3 and CD45) and 3 uropod markers (CD37, CD50, and CD162). Left in each column: 2D kamada-kawai graph layout of DNA-pixels A, each pixel colored in proportion to the log_10_(counts + 1) detected in the pixel. Right in each column: 3D density heat maps derived from the graph representations of the same cell colored by the count density of the indicated markers. The higher density areas are of yellow color, while low density areas are dark. F) Graph representation of Figure 4C, to emphasize the relationship between differentially colocalized proteins. Protein pair links are colored by the average difference in colocalization scores the pair displays in CD54 immobilized samples treated with RANTES, in comparison to control. The color scale is identical to Figure 4C.

These findings indicate that the polarization score and colocalization score show high concordance in detecting proteins with spatial rearrangement across conditions, while providing different yet complementary information about the spatial arrangement of proteins. While the polarization score indicates whether a protein is spatially clustered on the cell surface, the colocalization score reveals the protein’s spatial relationship with other proteins. A spatial pattern of proteins indicating a uropod signal is expected to exhibit both increased polarization and colocalization, as observed prominently in CD37, CD50, and CD162, all of which are known proteins located in the uropod. This exemplifies how MPX can be employed to identify patterns of protein spatial organization and their potential roles in cellular processes. In addition, the comparison of T cell immobilization effects on protein abundance (Figure 4A) and protein polarization (Figure 4B) revealed minimal overlap (Figure 5) with merely 2 proteins (CD11a and CD50) simultaneously showing significantly different abundance and polarization for the immobilized T cells treated with RANTES. These findings indicate a distinct and independent relationship between abundance and the two spatial metrics. Although we observed an effect on protein abundance, potentially associated with uropod formation, possibly characterized by reduced levels of uropod proteins CD43, CD50, and CD102, the interpretation of these changes is not straightforward. Notably, the uropod signal cannot be solely discerned from the abundance data, highlighting the importance of incorporating spatial metrics for a comprehensive understanding of protein behavior.

**Figure 5.**
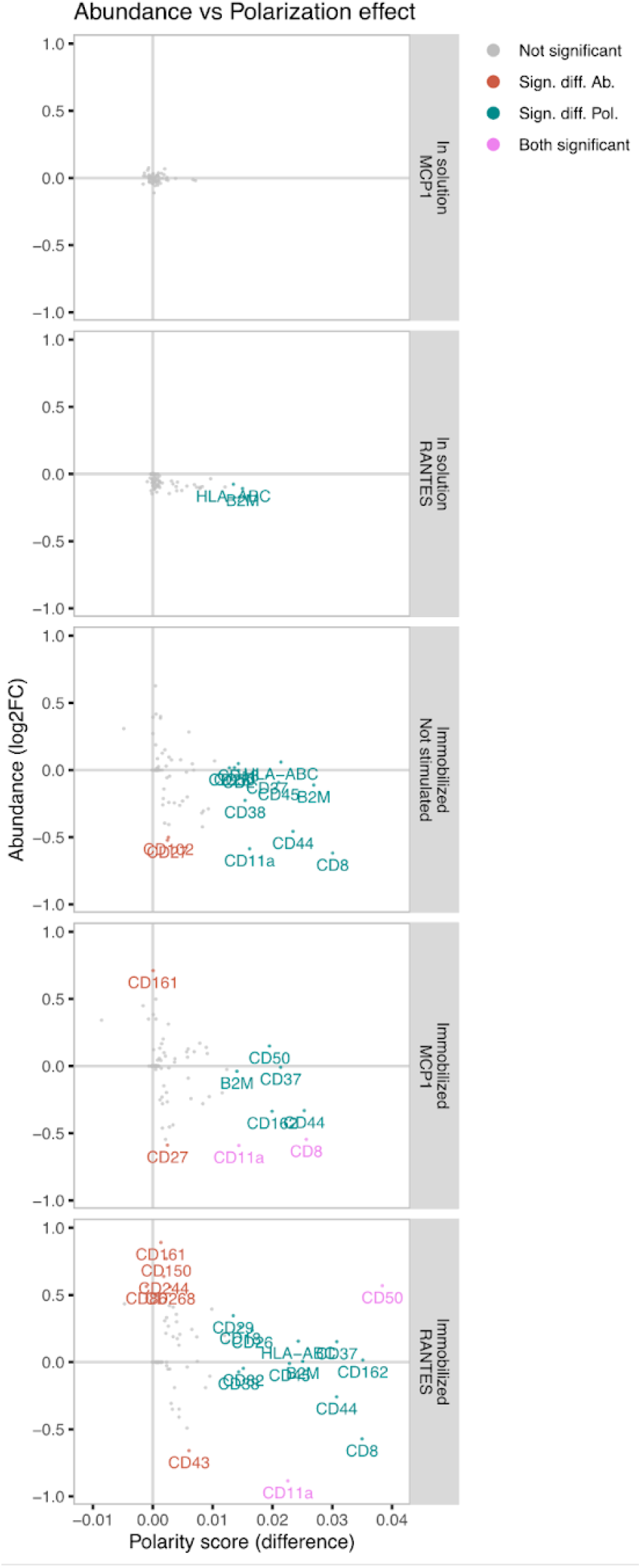
The effect of T cell immobilization and chemokine stimulation. Comparison of effect on target protein abundance (Y-axis) and polarization (X-axis) for each experimental condition compared to untreated.

## Summary

Converting proteins into DNA-tags to increase multiplexing, sensitivity and throughput has been a successful strategy to enable large scale targeted plasma proteomics studies (20). We anticipate that employing the DNA-tagging strategy also for targeted spatial proteomics will drive similar advancements. To the best of our knowledge, MPX is the first described method to identify relative localization of proteins in single cells without the use of microscopy. MPX offers a novel dimension to single cell analyses by revealing not only protein abundances, but also protein polarization and protein-protein colocalization patterns in single cells. These novel dimensions hold potential for new insights into essential cellular activities such as cell motility, cell activation, drug mode-of-action, and formation of cell-cell communication interfaces. Therapies that aim to reorganize cell surface proteins in order to support immuno-modulatory activities are currently under development (21, 22). The spatial arrangement of immune synapse proteins for chimeric antigen receptor (CAR) T-cell therapies can also be modulated to increase therapeutic efficacy by tuning CAR disorganized synapses into preformed synapses (23) illustrating the importance of single cell spatial proteomics beyond basic research.

The high level of multiplexing and throughput that MPX provides will enable discoveries in spatial proteomics. By representing cellular states as graph objects, it becomes possible to leverage spatial statistics metrics, including permutation testing to enable the identification and quantification of cellular changes relative to random expectations. Additionally, graph data is particularly well suited for neural networks and machine learning, which are increasingly influential in the realm of bioinformatics and computational biology (24). These features position MPX as a powerful tool for both method development and making new spatial proteomic discoveries.

The present work shows the applicability of MPX for single cell analyses as a multiplexed, 3D spatial proteomics method, without any dedicated instrumentation. As illustrated by the uropod formation study, high multiplexing with spatial dimensions can provide insights into T-cell motility, a vital process of tumor infiltration of lymphocytes and an essential process in immune therapies of cancer. MPX is anticipated to evolve to target other biomolecules and sample types, increased multiplexing, as well as benefit from the rapid advances in DNA sequencing, computational power, algorithm development, and machine learning.

## Methods

### PBMC extraction from whole-blood

PBMCs were separated from whole-blood collected in heparin or EDTA blood collection tubes by Ficoll-Paque density gradient centrifugation. The platelet fraction was reduced by 3 repeated centrifugation steps at 100 x g for 10 minutes.

### CD3 capping of PBMCs

PBMCs were first incubated with 50 μg/ml of human IgG for 15 min at 4°C to block Fc receptors and then incubated with 20 μg/ml of anti-CD3 AOC for 40 min at 4°C. After two washes, cells were incubated with 40 μg/ml of Goat anti-mouse secondary antibody for 40 min at 4°C, followed by incubation at 37°C for 1 hour. The capped cells were then immediately fixed with PFA.

### Rituximab stimulation of Raji cells

Raji cells were Fc-receptor blocked with 50 μg/ml of human IgG for 15 min at 4 °C and washed. Cells were then either PFA fixed directly or incubated with 20 μg/ml of Rituximab AOC in RPMI media for 60 min at 37 °C, followed by washing and PFA fixation.

### Uropod formation of T cells

PBMCs were first stimulated into PHA blasts with PHA-L for 48 hours, washed twice in PBS and incubated with 10 ng/ml of IL2 in RPMI media for 5 days at 37°C. Cell culture plates were coated with either 5 μg/ml of CD54Fc antibody alone or together with either 10 ng/ml of RANTES or MCP1 at 37°C for 1 h. Approximately 300,000 PHA blasts were added to each of the 3 coated plates for 2 hours. The adhered cells were then fixed with 1% PFA, before being brought back into suspension via incubation with TrypLE enzyme solution (ThermoFisher) for 10 min at 37°C. For the solution conditions, PHA blasts were instead incubated in solution with either 10 ng/ml of Rantes or MCP1, followed by PFA fixation.

### Cell fixation and AOC staining

Cells resuspended in PBS were fixed in for 15 min at room temperature in a fixation solution containing 1% v/v PFA in PBS. Cells were washed once in PBS, followed by addition of a Blocking/Quenching buffer containing 1% FBS, 0.1% BSA, 1 mg/ml ssDNA, 50 μg/ml human IgG, 125 mM Glycine, 4 mM EDTA, 0.04% Proclin300 in PBS. Cells were incubated for 15 min at 4°C, followed by a wash in PBS to remove the blocking/quenching solution.

Fixated and blocked cells were stained for 30 min at 4C in a 50ul reaction containing a cocktail of 80 AOCs, each at a concentration of 5μg/ml, in a staining buffer comprising 0.2% BSA and 2 mM EDTA in PBS. After 3 washing steps in Wash Buffer (50 mM NaCl, 1 mM EDTA, 20 mM Tris-HCl, pH8), AOCs bound to cells were stabilized using a secondary antibody by incubating the cells for 30 min at 37°C in a secondary antibody solution consisting of 20 μg/ml secondary antibody (25), 0.2% BSA and 2 mM EDTA in PBS, followed by two washing steps in Wash Buffer, before proceeding with MPX protocol.

### Molecular Pixelation protocol

DNA pixels were allowed to hybridize to AOCs on 15 000 stained cells in a 50 μl A-pixel hybridization reaction containing 1 nM A-pixels, 2 μM A_GFP oligo, 1 μg/ul of sheared Salmon sperm DNA, in a hybridization buffer containing 300 mM NaCl, 15 mM MgCl2, 20 mM Tris-HCl (pH 8) and 0.05% Tween20. The reaction was incubated for 15 min at 55°C, followed by 2 washes in Wash Buffer.

A-pixel unique pixel identifiers (UPI) were incorporated onto the mab-oligos via a 50 μl gap-fill ligation reaction consisting of 40 U Taq ligase (New England Biolabds) 3 U T4 DNA polymerase (New England Biolabs), 100 μM dNTPs, 0.5 mM NAD^+^ in 1x rCutSmart buffer (New England Biolabs). The gap-fill reaction was incubated at 37°C for 20 min, followed by 1 wash in Wash Buffer.

A-pixels were degraded by incubating the cells in a 50 μl reaction containing 1 U USER enzyme (New England Biolabs) in Wash Buffer. The reaction was incubated at 37°C for 30 min, followed by 1 wash in Wash Buffer.

Hybridization of B-pixels was performed in the same reaction conditions as described for A-pixels with the exception of using B_GFP oligo instead of A_GFP oligo. Similarly, the gap-fill ligation reaction for B-pixels was performed in the same reaction conditions as described for A-pixels.

In order to remove any incomplete assay products, an exonuclease treatment targeting DNA with unprotected 5’ ends was performed. Cells were first counted using a hemocytometer and an aliquot of less than 1000 cells were resuspended into a 15 μl reaction containing 10 U Lambda exonuclease, 100 μM dNTPs, 1 mM NAD+ in 1x rCutSmart buffer (New England Biolabs). The reaction was incubated at 37°C for 30 min, followed by heat inactivation at 75°C for 10 min.

### PCR & NGS sequencing

PCR was performed in a 40 μl PCR reaction containing 1x Q5 HotStart Hifi PCR master mix (New England Biolabs), 0.4 μM of Illumina adapter PCR primers (ILM_p5_PCR, ILM_p7_PCR) containing 8 nt sample indexes to allow multiplexing and 15 μl of sample from the lambda exonuclease step.

The PCR products were purified twice using AmpureXP SPRI beads (Beckman-Coulter) according to manufacturer’s instructions and subsequently quantified using Qubit HsDNA assay (ThermoFischer). The purified PCR products were diluted to 0.65 nM with 15% phiX spiked in and paired-end sequenced on an Illumina NextSeq2000 sequencing system, using 44 cycles for read1 and 78 cycles for read2.

### Antibody oligonucleotide conjugates (AOC) preparation

Monoclonal antibody clones were first validated one-by-one for specific target binding on PFA fixed PBMC. Then these were coupled to oligonucleotides by DBCO-Azide click-chemistry (26), Supplementary Table 1 and Table 2.

### RCA template preparation

Circularized DNA templates were prepared by incubating 300 nM of template oligo with 200 nM padlock probe in a 50 μl ligation reaction containing 1mM ATP and 400U of T4 DNA ligase (New England Biolabs) in a buffer comprising 33 mM Tris.acetate (pH 7.9), 10 mM Magnesium acetate, 66 mM potassium acetate, 0.1% Tween20 and 1 mM DTT. The reaction was incubated for 30 min at 37°C, followed by heat inactivation at 75°C for 10 min. To each ligation reaction, 10 U of Exonuclease I and 20U of Exonuclease III (New England Biolabs) was added to each reaction, and incubated at 37°C for 30 min, followed by heat inactivation at 85°C for 20 min. See Supplementary Table 2 for oligonucleotide sequences.

### DNA-pixel preparation

DNA pixels were prepared in 75 μl RCA reactions comprising 5 nM circularized RCA template, U phi29 enzyme (ThermoFisher), 0.75 mM dAUGC or dNTPS, in the same reaction buffer as used for RCA template preparation. The reactions were incubated at 37°C for 4 min, followed by heat inactivation at 65°C for 10 minutes. 1U of rSAP enzyme was added to each reaction in order to inactivate free dAUGCs or dNTPS and incubated at 37°C for 20 min, followed by 5 min at 65°C. See Supplementary Table 2 for oligonucleotide sequences. DNA-pixels were analyzed by electron microscopy and shown to be spherical molecules ranging between 50 and 250 nm, Supplementary Figure 10. Assuming they cover the entire cell surface area of a typical lymphocyte at 92 um^2 (27), the average DNA-pixel diameter is estimated at around 200 nm.

### Data analysis by Pixelator

MPX sequencing data was analyzed by a dedicated MPX data analysis pipeline, Pixelator. First, sequence reads were quality filtered to remove low quality reads. Next, reads were matched against the common pixel binding sequence motifs (BS1 and BS2) and reads with > 10% mismatch were discarded. The BS1 and BS2 sequences were then discarded so that only the UMI, BC, UPIA and UPIB sequence motifs were kept for each read. Duplicate reads generated from the PCR step prior to sequencing were collapsed into unique sequences defined by the combination of the 10nt UMI and the 25nt UPIA sequences. Correction of PCR and sequencing errors were performed by clustering the set of putative unique reads based on hamming distance and identifying a consensus sequence from read counts from those sequences that clustered together. The UPIA, UPIB, UMI and BC sequence motifs were extracted from each unique read and stored as an edge list. An undirected graph was generated from the UPIA and UPIB sequences of the edgelist from a sequenced sample.

Community detection based on modularity maximization was performed on the resulting graph in order to identify and remove any spurious edges connecting densely connected communities of the graph assumed to represent cells, followed by inducing subgraphs for each connected component of the graph. Subgraph memberships were assigned to each edge of the edgelist and an AOC count matrix was generated by summing up counts for each protein for each subgraph membership of the edgelist.

From the total AOC count (number of edges) of each subgraph membership, size outliers were identified based on the descending rank order distribution from total AOC count values per subgraph. A size threshold based on the rate of change was defined by finding the first and second derivatives from an univariate smooth spline curve fitted to the linear-log distribution of the ranked antibody count data. Edges corresponding to subgraph memberships considered as size outliers were removed from the edgelist.

Finally, the filtered edgelist was used for downstream analysis such as calculation of Polarity scores and colocalization scores by interrogating the graph constituting each cell. The graph generated from the UPIA and UPIB sequences of the edge list is bipartite, forming only edges between UPIA and UPIB nodes, and never between two UPIAs or two UPIBs. The one-mode projections of the graphs, with edges linking directly between UPIA nodes, and the set of antibody counts associated with each UPIA node as node attributes, were used for the purpose of calculating spatial metrics such as Polarity scores and colocalization scores.

### Polarity score calculation

Spatial autocorrelation was used to quantify the degree of clustering or non-randomness of the spatial distribution of a protein within a cell graph. Moran’s Index (I) of spatial autocorrelation were computed from spatial weights defined by the row-normalized adjacency matrix of the one-mode projected cell graphs. In addition to the Moran’s I value, a z-score and p-value was computed under the randomization null hypothesis for each cell graph. The obtained z-scores thus denoted the degree of non-randomness, or clustering, of the protein in the context of the cell graph it was calculated from, and was used as Polarity scores.

### Colocalization score calculation

The calculation of colocalization score consisted of 5 steps, which were performed for each individual component, cell. 1) Count filtering, which removed data points that had too little data to generate a robust score, disregarding markers that have too few counts, or spatial locations (UPIAs) which have too few counts. 2) Permutation, which generated simulated components with permuted protein localization that were used to create a null distribution that enabled the calculation of how much the resulting score deviated from random colocalization. 3) Scaling and transformation, which applied normalizations to the count data of the original component as well as the permuted components. 4) Calculation of the colocalization statistic; Pearson’s r. 5) Null distribution fitting, which compares the observed score to the scores of the permuted components to generate a p-value and Z-score that describe the statistical significance of the score, i.e. the degree to which the score was lower or higher than what was expected by chance.

### Graph visualization in 3-D

Three approaches were used to generate graph layouts for visualization. In the first approach euclidean coordinates in three dimensions were generated for each node by applying the force-directed graph layout algorithm kamada-kawai on the graph representation of a cell. For the second approach, the generated coordinates were projected onto the unit sphere by dividing each coordinate by its norm. The subset of graph nodes associated with a protein marker of interest were colored.

For the third approach, a density heatmap was generated from the sphere projected coordinates. A color value representing count density was obtained for each point of a three dimensional grid representing the surface of the unit sphere by applying a function which iterated over each surface grid point and calculated the distance, defined by the euclidean norm, to each of the sphere projected node coordinates associated with a specific protein target. For all node coordinates within a selected distance cutoff value to each grid point, 1 - distance / distance_cutoff were summed up and the logarithm of this sum was used as the color value for each grid point.

### PBMC data analysis

The cells were filtered to remove size outliers, removing the 10 largest cells from each replicate, and cells with fewer than 4,000 detected UMIs (Supplementary Figure S11). Next, the cells were filtered for outliers in terms of distribution of counts across AOCs using the Tau metric (28), a numerical value between 0 and 1 describing the degree to which skewness the distribution of counts across markers. Evenly distributed across markers generates a Tau of 0, and a distribution completely skewed to a single marker results in a Tau of 1 (See methods). Antibody aggregates will often consist of a random composition of antibodies (high complexity) or are composed of a single antibody (low complexity), resulting in a notably low and high Tau score respectively (Supplementary Figure S12). Across these two samples, a single high complexity aggregate was identified and removed. After applying cell size cutoffs and aggregate removal, 388 and 495 cells remained, respectively.

### CD3 data analysis

The data output from 3 separate experiments containing CD3-capped and control samples were combined and analyzed in aggregate. The protein count matrix output from each sample was centered-log-ratio (clr) transformed across proteins and filtered to only retain the T-cell fraction by applying cutoffs for T cell markers of 1.6 for CD3, and >2.5 for either CD4 or CD8. Any remaining non T-cells were removed from the data by filtering the cell-count matrix for CD19, CD14 and CD20 clr-transformed counts being below 1, 1 and 2 respectively. The remaining T-cells were subsequently filtered based on total total antibody count below 15 000 to remove size outliers. After the filtering steps, 2,075 CD3-capped and 1,359 control cells remained. Polarity scores for CD3 were compared between control and stimulated conditions and a p-value calculated using the Wilcoxon Rank Sum test.

### CD20 data analysis

The data output for each cell was size-filtered on total antibody counts to remove size outliers to retain cells with UMI counts between 15 000 - 60 000 UMIs. The remaining cells were filtered to remove any cells with clr-transformed counts for CD19 < 2. Polarity scores were calculated for the remaining 597 stimulated cells and 442 control cells, and a p-value for CD20 (Rituximab) Polarity scores between control and stimulated samples were calculated using Wilcoxon Rank Sum test.

## Data availability

(not available in bioRxiv preprint)

## Code availability

(not available in bioRxiv preprint)

## Acknowledgements

The presented work was funded by the Wellcome Leap ΔTissue Program and Stiftelsen för Strategisk Forskning, SSF.

## Author information

### Author affiliations

All authors at Pixelgen Technologies AB, Nanna Svartz väg 2, 17165 Solna, Sweden, except Petter Brodin

Simon Fredriksson, Department of Protein Science, Royal Institute of Technology, SciLifeLab, Stockholm, Sweden.

Petter Brodin, Dept. of Women’s and Children’s Health, Karolinska Institutet, 17165, Solna, Sweden and the Dept. of Immunology and Inflammation, Imperial College London, W12 0NN, London, UK

### Contributions

FK and SF conceived of the Molecular Pixelation method. FK, SF, TK, DT, LL, SG, MK, EP, JB, MGG, JRK, JD, MS, and PS designed and/or performed experiments validating the methods and developed the specific reagents needed. FK, AMB, JFN, JD, MK, JRK developed the data analysis algorithms and the pipeline. FK, MK, AMB, PB, DT performed in depth analyses of biological results. SF, FK, MK, PB, and AMB wrote the manuscript. All the authors read and approved the manuscript before submission.

## Ethics declaration

All PBMC samples were purchased from a Karolinska Hospital blood bank drawn from healthy volunteers with informed consent and withheld sample identity or other medical information.

