## Supplementary information for "Molecular Pixelation: Single cell spatial proteomics by sequencing"

Figure S1 - Scatter plot of cell input linearity
Figure S2 - Heatmap of count levels for PBMC sample
Figure S3 - Violin plot of Isotype and intracellular control count levels
Figure S4 - Histograms of AOC counts, UPI counts, read counts for PBMC sample
Figure S5 - Volcano plots for polarity scores FOr CD3 capping and rituximab stimulation
Figure S6 - Polarity scores CD3 capping & rituximab additional targets
Figure S7 - Graph layouts of representative Uropod cells
Figure S8 - Three graph layout approach examples for Uropod cell
Figure S9 - Pairwise Colocalization patterns represented as weighted graphs
Figure S10 - Electron microscopy image of DNA pixels
Figure S11 - Edge rank plot of AOC counts per cell for PBMC sample
Figure S12 - Scatter plot of Tau scores for PBMC sample

Table S1 - Antibody clones
Table S2 - Oligonucleotide sequences

**Supplementary animations** (not available in bioRxiv preprint)
CD3-capped T cell.mp4
Uropod_RCVCMP0000275_CD50_anim.mp4
Uropod_RCVCMP0000275_CD162_anim.mp4
Uropod_RCVCMP0000275_CD45_anim.mp4

**Supplementary Figure S1**. Cell input range linearity.

One sample was processed with Molecular pixelation and cells were counted prior to PCR reaction. 200, 500 or 1000 cells were input to separate PCR reactions, and carried forward to sequencing. The number of detected cells after processing the data through the data analysis pipeline correlated well with the number of input cells into PCR ($R^2$ value of 0.99). The number of detected cells were 161, 517 and 1054 for the inputs of 200, 500 and 1000 cells respectively. Variability from counting, dilution and aliquoting cells into PCR likely contributed to some of the deviation from expected input.

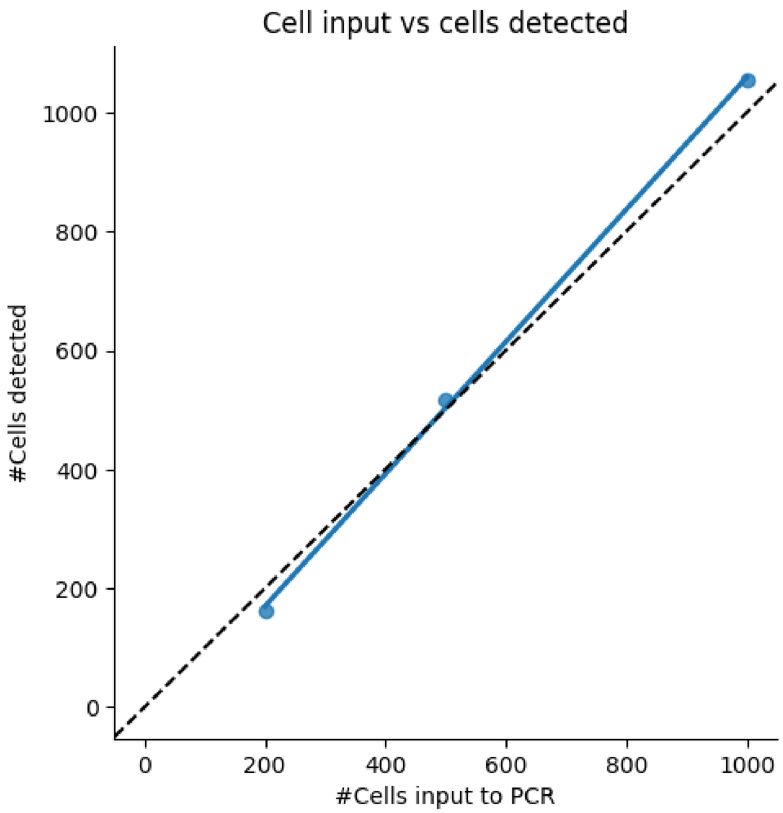

**Supplementary Figure S2.**

Heatmap of scaled counts of all 80 markers (x-axis; including the 4 control markers) across all cells after filtering (y-axis; n = 814). The two tracks to the left of the heatmap denotes the source sample and the cell type annotation of the components. The color corresponds to relative counts multiplied by a factor of 10000, and then log transformed (log10(x + 1)).

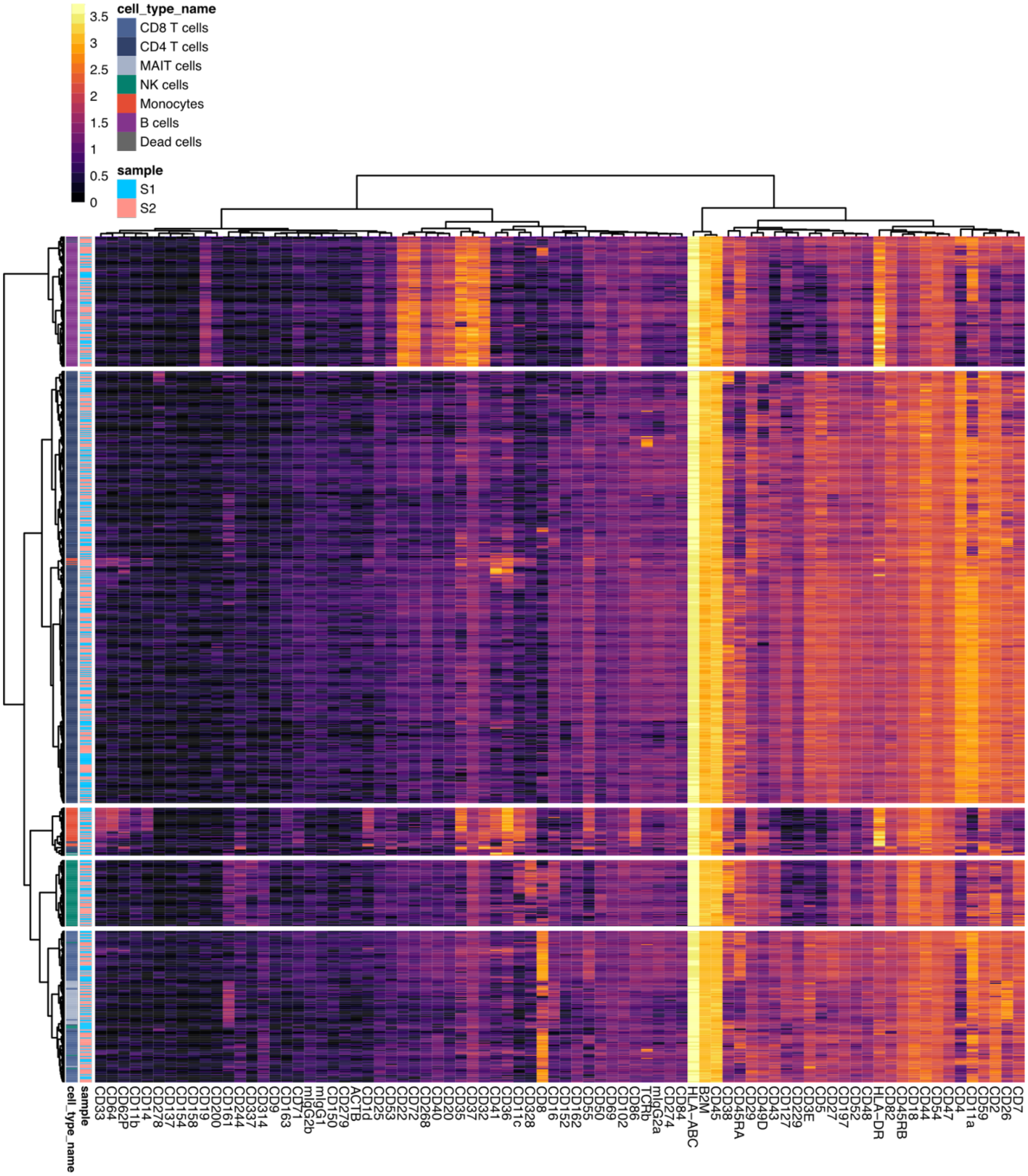

**Supplementary Figure S3.**

Violin plot showing the distribution of relative counts (%) for isotype markers in unstimulated cells. The median percentage is marked.

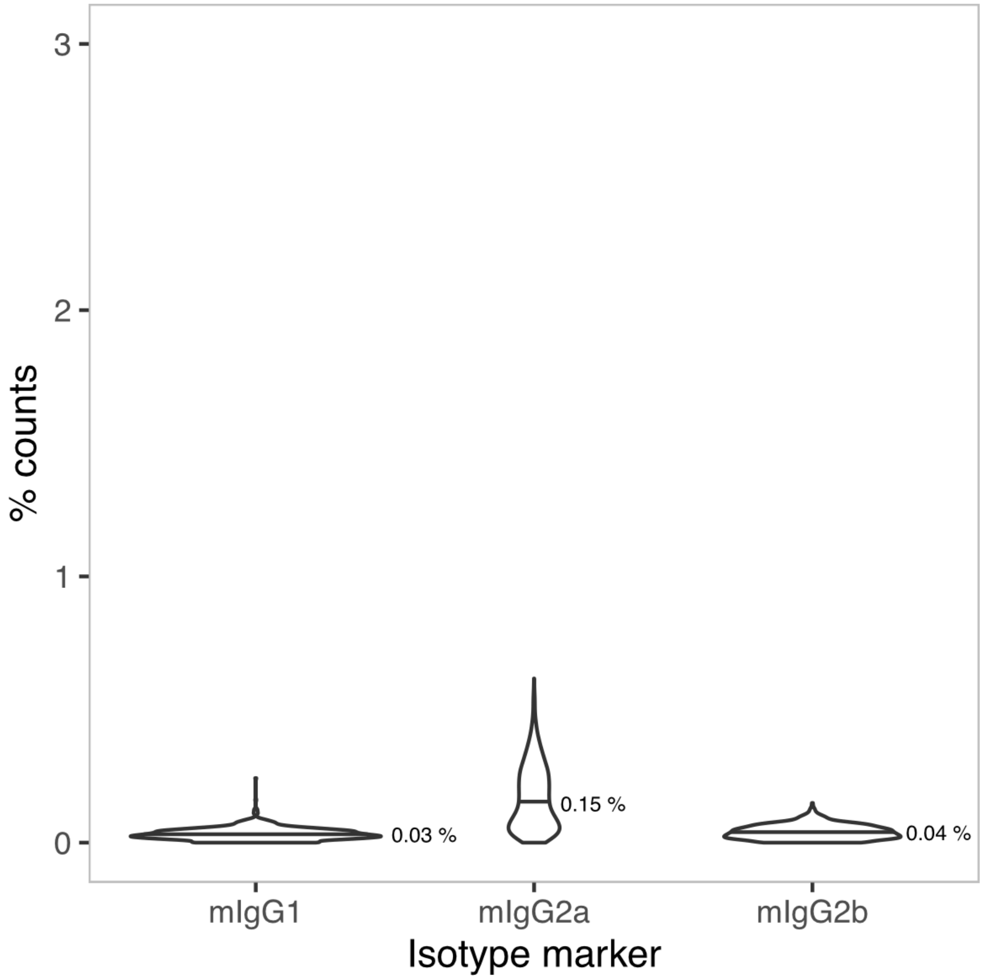

**Supplementary Figure S4.**

Distribution of metrics per component (single cell). Histograms showing the distribution of the number of reads, edges (UMIs), DNA-pixels A (UPIA), DNA-pixels B (UPIB), and the number of unique molecules per DNA-pixel A (UMIs/UPIA). The median of each metric is marked by a dashed line.

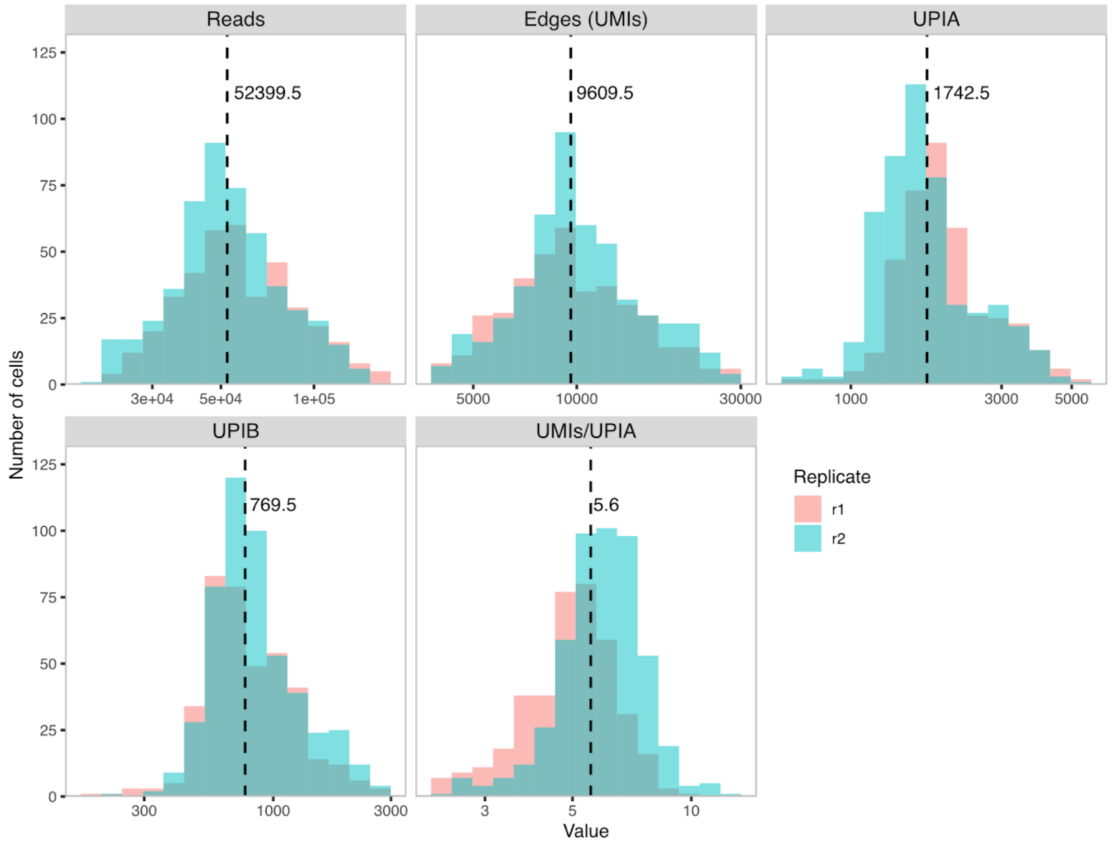

**Supplementary Figure S5.**

Volcano plots for the differential polarization analysis (Wilcoxon Rank Sum test) for A) T cells in PBMC stimulated by CD3 capping and B) Raji cells stimulated with Rituximab.

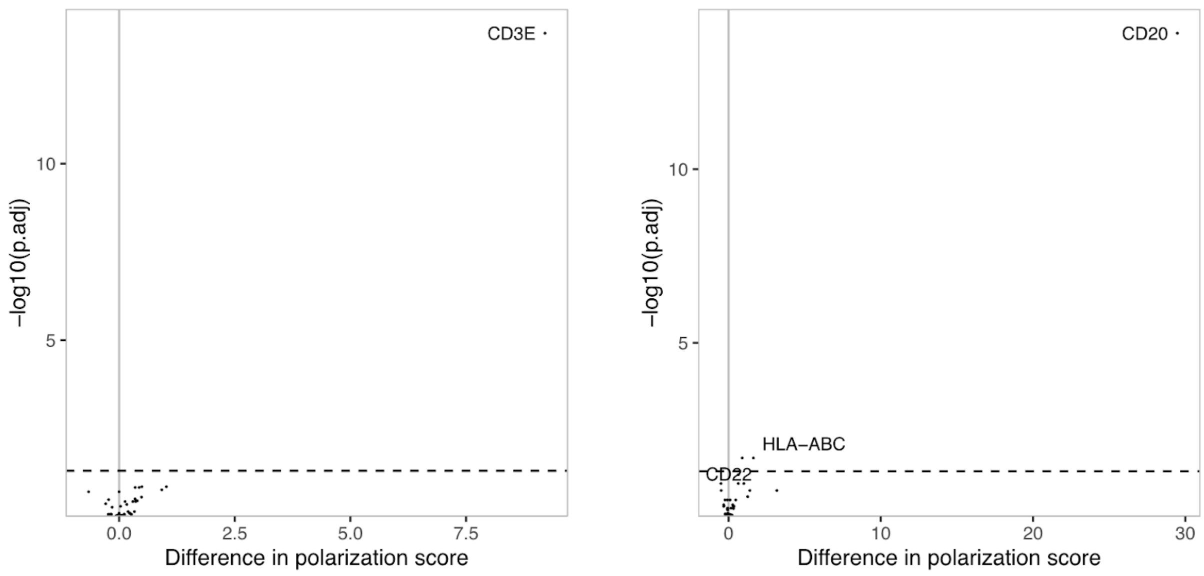

**Supplementary Figure S6.** Polarity scores for stimulated and control conditions of all markers with total counts above isotype control levels for the CD3 capping experiment (A) and Rituximab treatment experiment (B)

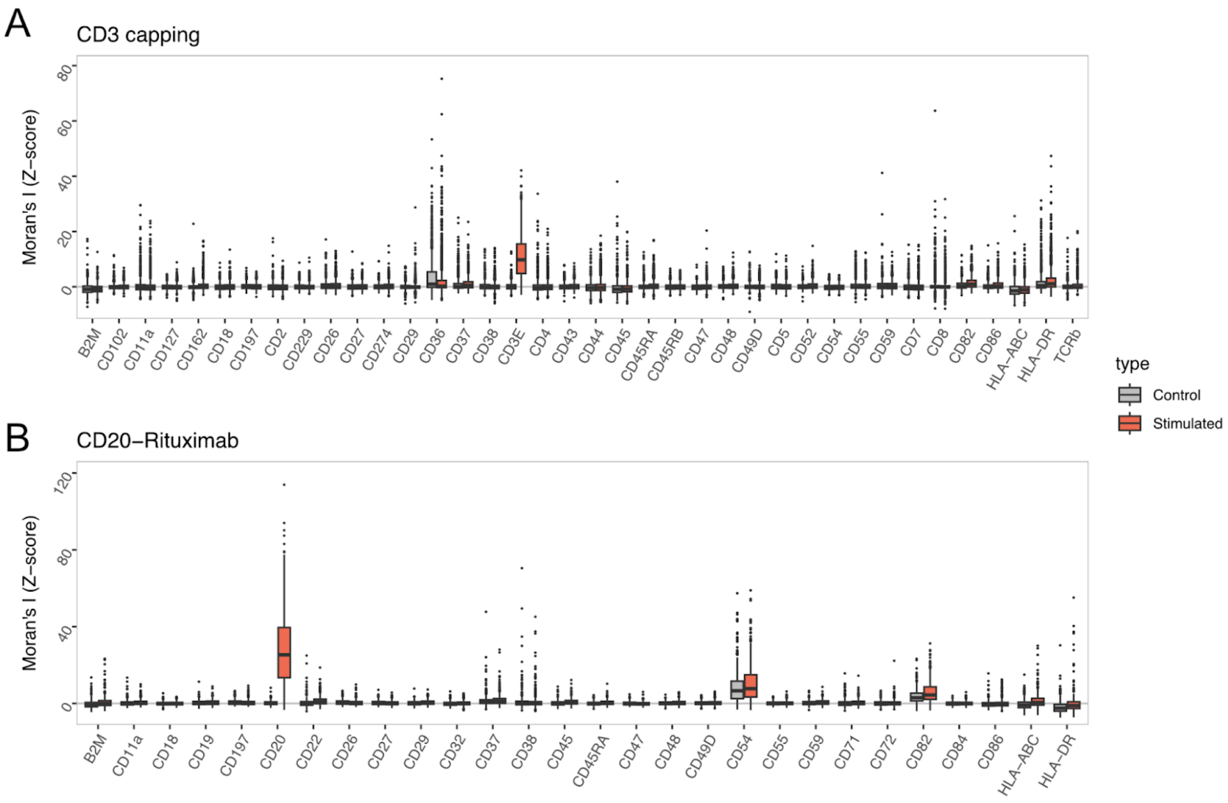

**Supplementary Figure S7.**

Graph visualizations generated by the kamada-kawai graph layout algorithm of single cells displaying colocalization of uropod markers. Two ubiquitous “background” markers and 3 uropod markers are shown in 9 cells from the sample exposed to the CD54 coated surface and stimulated with RANTES. Each point is an individual DNA-pixel A, and its color is proportional to the log-scaled expression of the marker. The coordinates of the DNA-pixels were generated by applying the force-directed graph layout algorithm kamada-kawai on the bipartite graph representation of each cell.

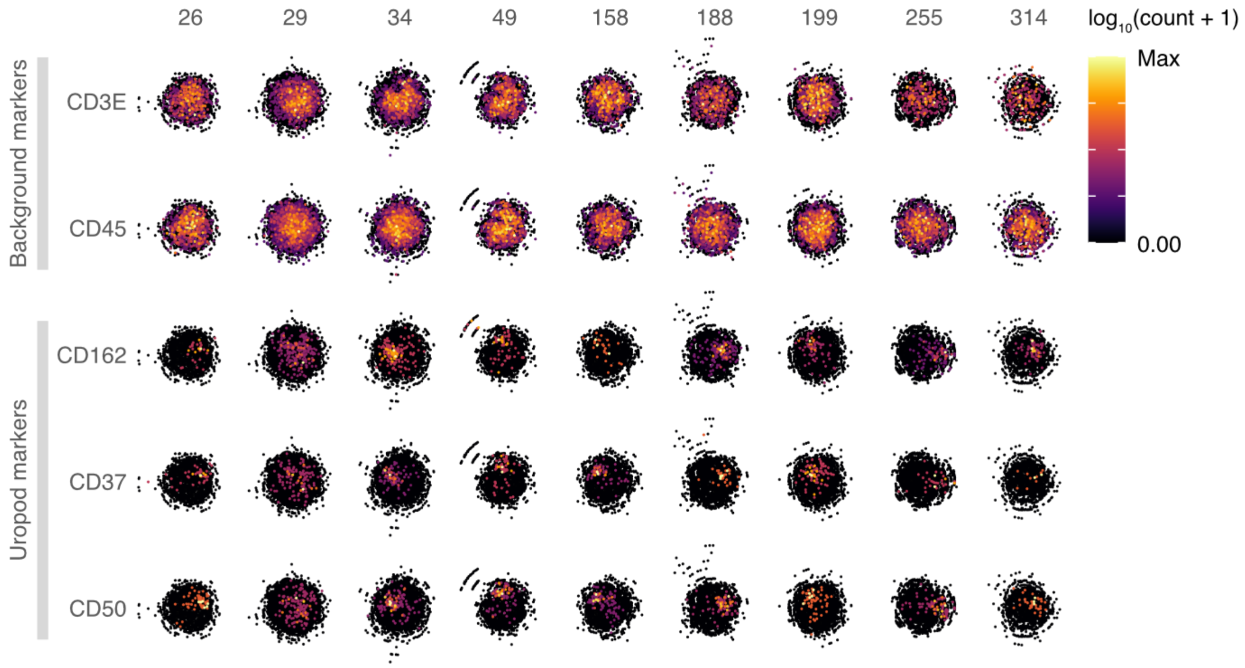

**Supplementary Figure S8.**

Examples of 3 approaches for 3D graph visualization layouts for proteins CD50, CD162 and CD3 from one selected chemokine-stimulated cell. Left: Layout obtained from force directed layout algorithm kamada kawai (kk). Middle: Projection of coordinates from kk layout onto unit sphere. Right: Heat map interpolation of count density onto a sphere surface. See methods for further details. The plots show a clustered or polarized protein arrangement of CD50 and CD162 in the same location, thus also showing colocalization of these markers, while CD3 is randomly distributed across the cell graph.

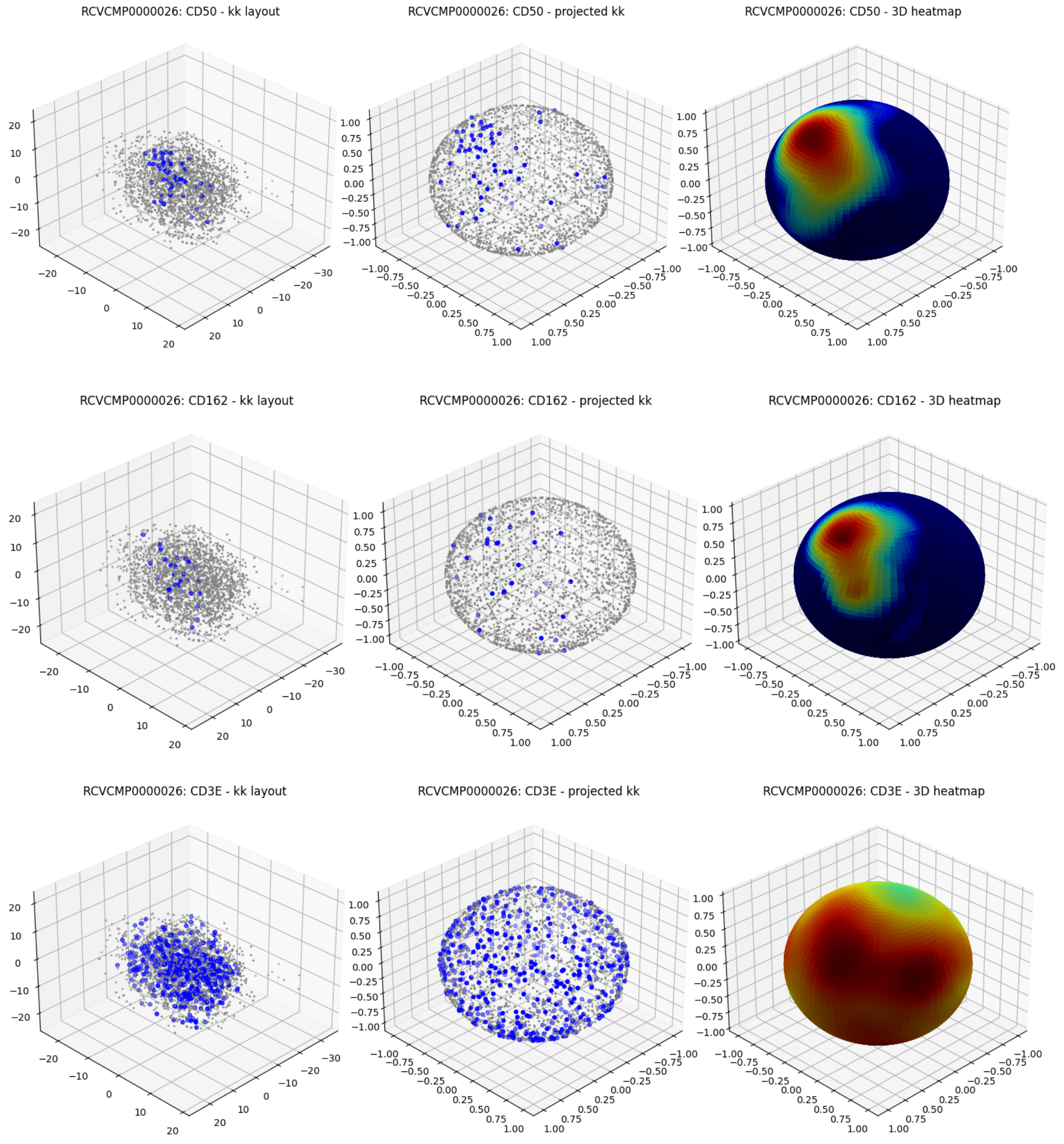

**Supplementary Figure S9.**

Network visualizations of protein pairs that are differentially colocalized across any of the 5 conditions (CD54 coating and/or chemokine stimulation) compared to the in solution unstimulated control sample. The difference (increase or decrease) in colocalization between a pair of proteins is shown as the color of the link (blue = decrease, red = increase), and both the size of the link and its color intensity shows the magnitude of the difference. The level of the Benjamini Hochberg adjusted p-value is denoted as follows: '*' = p < 0.05, '**' = p < 0.01, '***' = p < 0.001, '****' = p < 0.0001. Colocalization differences that are not statistically significant are still shown, but without the p-value annotation.

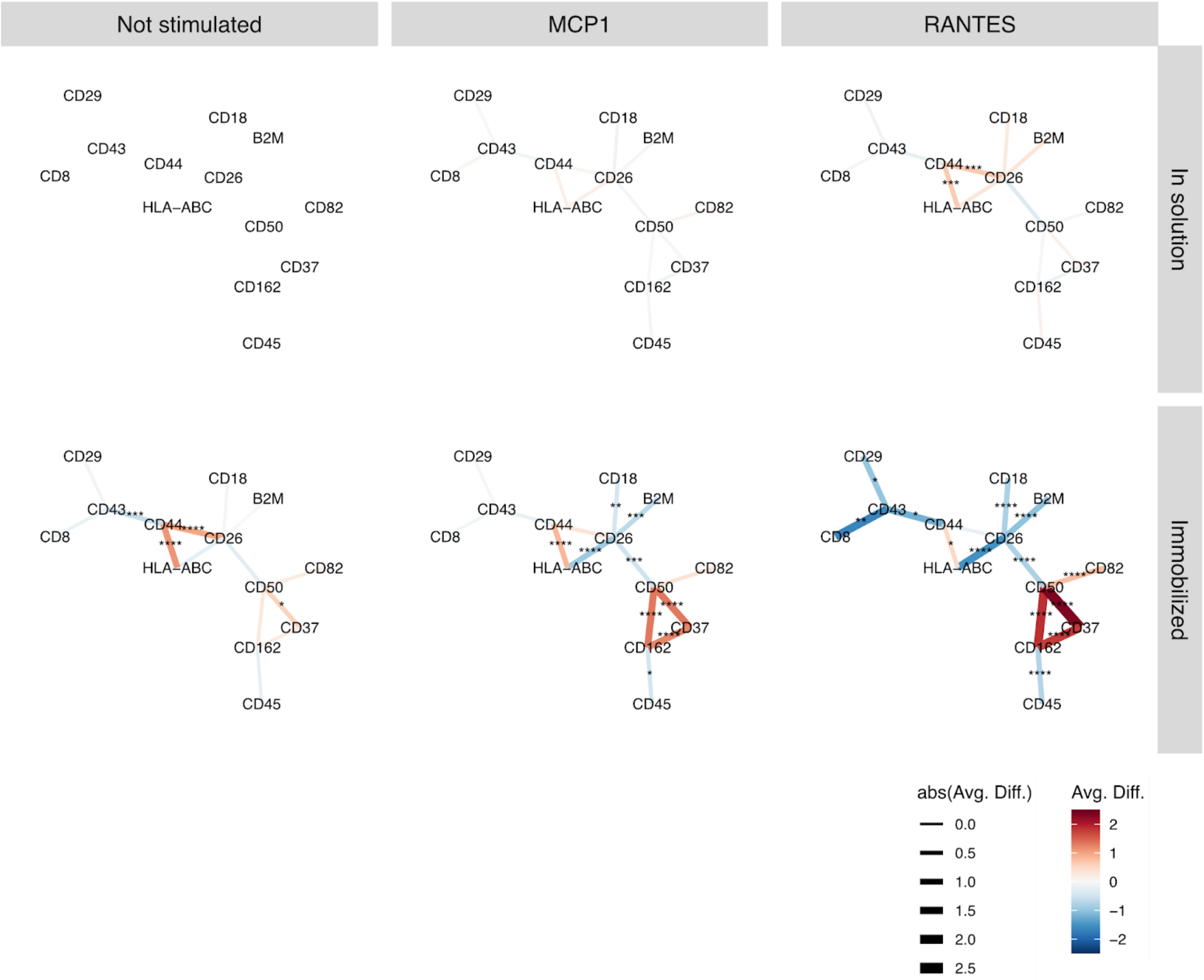

**Supplementary Figure S10**. Scanning Electron Microscopy (SEM) of DNA-pixels. SEM imaging performed by Swedish National Microscopy Infrastructure, NMI (VR-RFI 2016-00968). RCA-products collapse into spheres which has previously been shown by SEM (Deng et al DNA-Sequence-Encoded Rolling Circle Amplicon for Single-Cell RNA Imaging. Chem 4, 1373–1386, June 14, 2018)

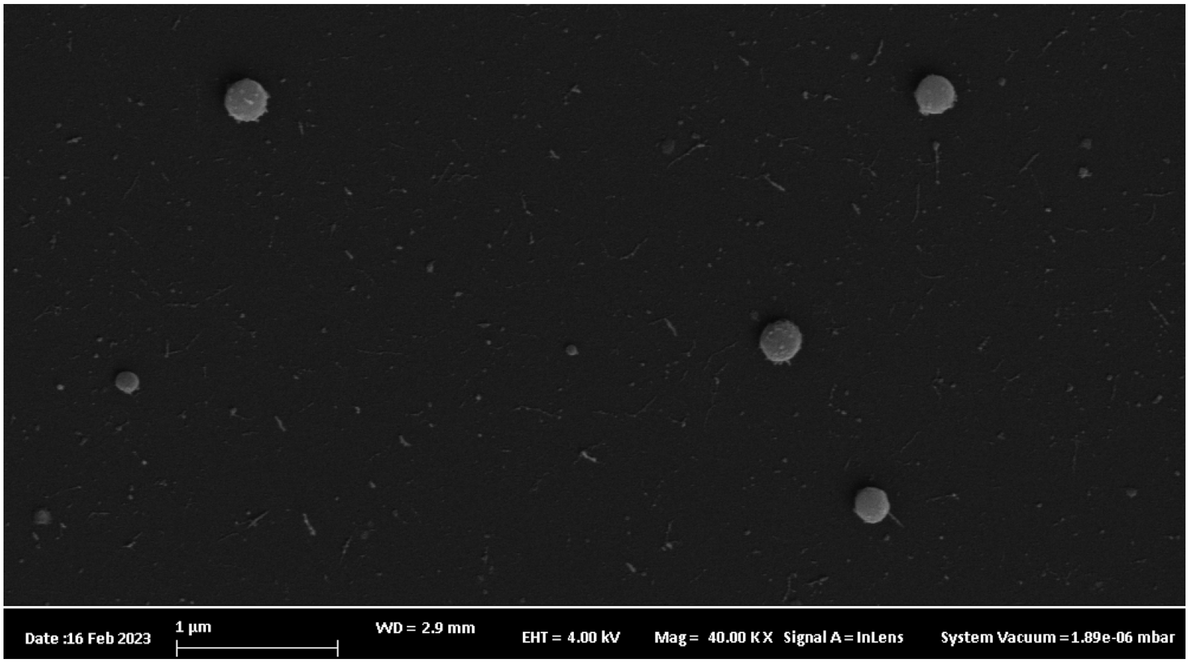

**Supplementary Figure S11.**

Edge rank plot for the two replicates of healthy PBMC, analogous to the “Barcode Rank Plot” often used to visualize cell calling in scRNAseq. The y-axis displays the number of edges (UMIs) per cell, which are ranked from highest to lowest along the x-axis. The two dashed lines mark the hard cutoffs for filtering the smallest and largest cells.

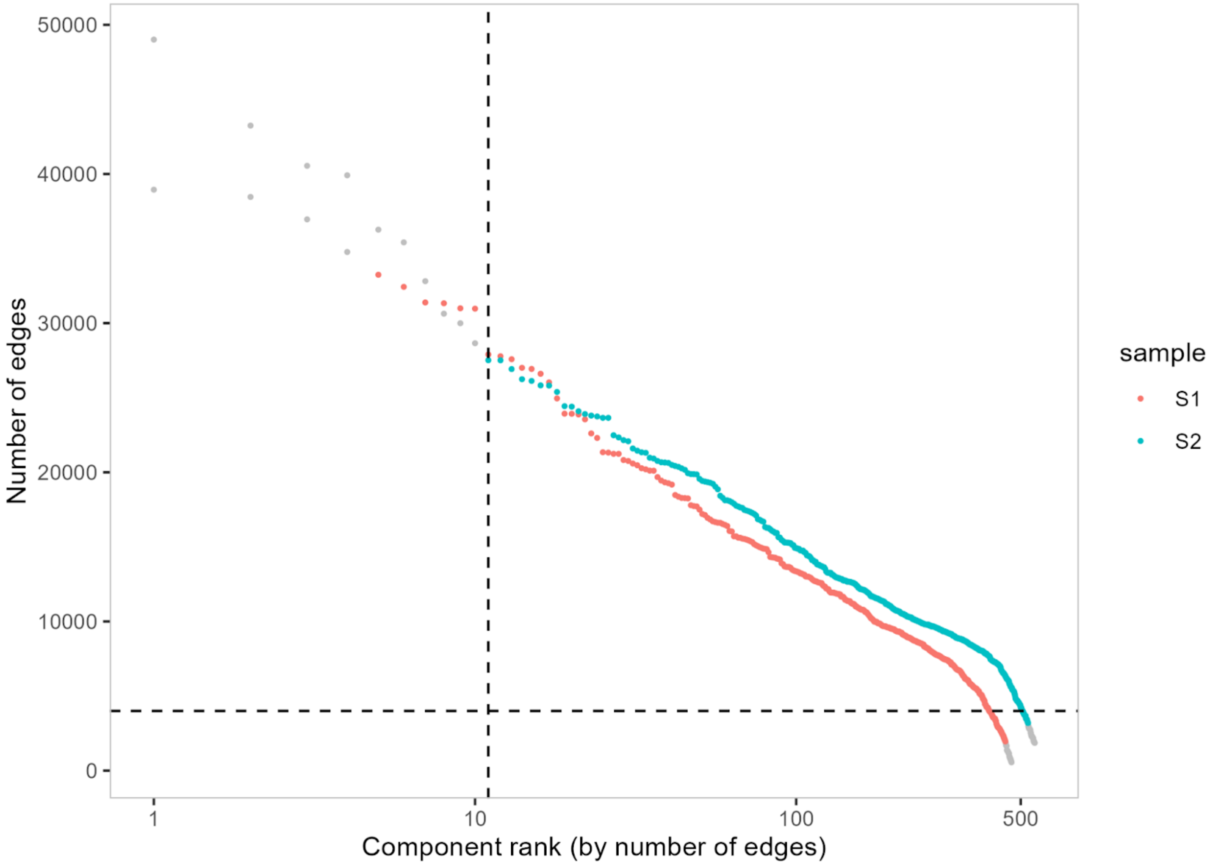

**Supplementary Figure S12.**

The DNA-pixel content (UMI/UPIA) versus the marker specificity (Tau score) for cells in the healthy PBMC samples. A single outlier, identified as a potential antibody aggregate was identified in sample 2 (marked in teal).

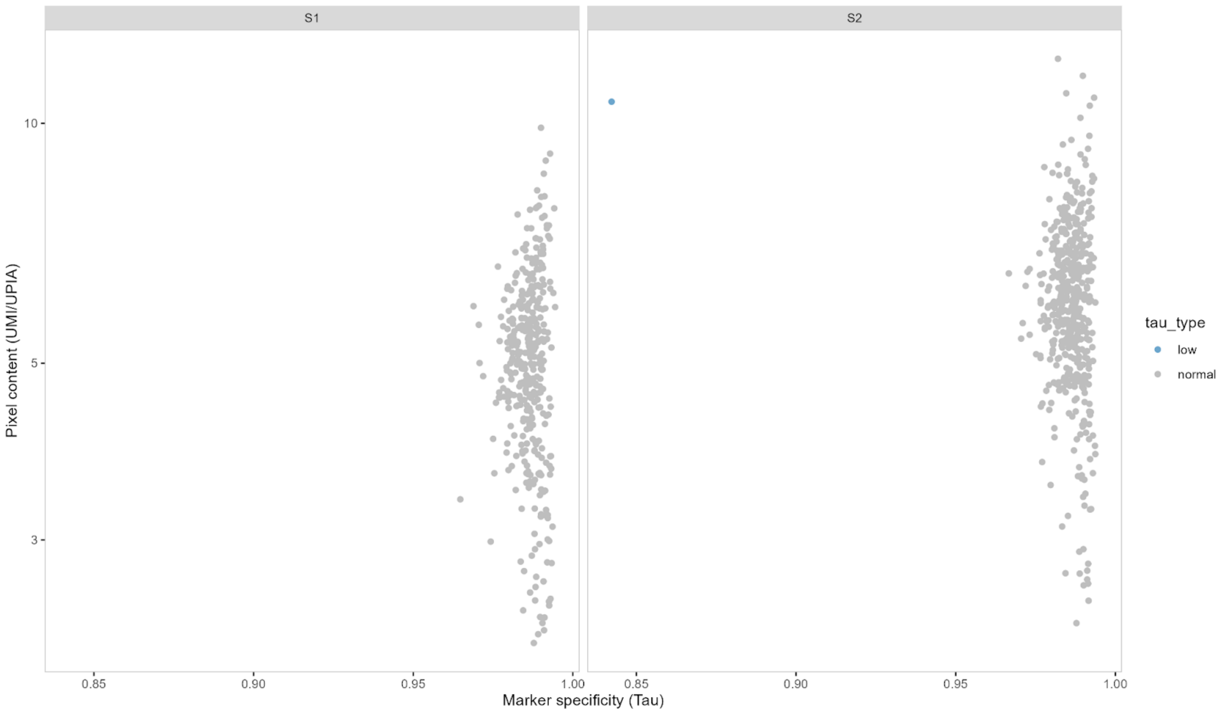

**Supplementary Table 1**. Monoclonal antibody clones and barcodes.

| Target name | Clone | Barcode name | Barcode sequence |
|---|---|---|---|
| B2M | B2M-02 | BC48 | CTGTAGGA |
| BAFFR | 11C1 | BC104 | CACGTTTC |
| CD102 | CBR-IC2-2 | BC84 | TTTCTGGT |
| CD11a | HI111 | BC18 | ACATTGAC |
| CD11b | ICRF44 | BC32 | ACTGTGTC |
| CD11c | Bu15 | BC17 | AAGTCGTG |
| CD127 | A019D5 | BC19 | TGATAGAA |
| CD137 | 4B4-1 | BC61 | CCTAAGAA |
| CD14 | 61D3 | BC40 | AGAGGCTC |
| CD150 | A12 (7D4) | BC72 | CTTGCACC |
| CD152 | BNI3 | BC51 | AACGGCTA |
| CD154 | 24-31 | BC64 | TGGGGCTT |
| CD158 | NKVFS1 | BC108 | ACTCGGAA |
| CD16 | 3G8 | BC14 | GTGCATTC |
| CD161 | HP-3G10 | BC57 | GTGAGTAG |
| CD162 | KPL-1 | BC80 | GCTATTGA |
| CD163 | GHI-61 | BC58 | CATGGGCA |
| CD18 | TS1-18 | BC65 | CACACGGT |
| CD19 | SJ25-C1 | BC26 | CTACGACG |
| CD197 | G043H7 | BC34 | AGGATGTT |
| CD1d | 51-1 | BC54 | TACTCTTG |
| CD2 | TS1-8 | BC37 | CCGATATC |
| CD20 | 2H7 | BC38 | GAATCCCG |
| CD200 | OX-104 | BC59 | AGGGCAGT |
| CD22 | S-HCL-1 | BC73 | CTCAAGAG |
| CD229 | HLy9-25 | BC62 | GTTCAGAC |
| CD244 | C1.7 | BC63 | AGCCCGAA |
| CD25 | BC96 | BC4 | GCCGGACG |
| CD26 | BA5b | BC11 | GTCTTTGT |
| CD27 | LT27 | BC12 | GTTGTCCG |
| CD274 | 2A3 | BC1 | TCCCTTGC |
| CD278 | C398-4A | BC101 | AAAGCAAC |
| CD279 | EH12-2H7 | BC5 | TTCTGGGT |
| CD29 | 3B6 | BC20 | ATTCGCCT |
| CD314 | 1D11 | BC109 | CTTCTTGA |
| CD32 | FUN2 | BC102 | CAATCGGC |
| CD33 | WM53 | BC77 | TCCGTAAC |
| CD337 | P30-15 | BC53 | TCCCAGTG |
| CD35 | E11 | BC110 | CCAGACAC |
| CD36 | 5-271 | BC49 | ATTTCGAG |
| CD37 | IPO-24 | BC79 | TTGTCCAA |
| CD38 | HIT2 | BC13 | TCAACGCT |
| CD3E | UCHT1 | BC36 | AGCTACTA |
| CD4 | OKT4 | BC42 | CTGACCAT |
| CD40 | C40-1605 | BC25 | TCAGGGTG |
| CD41 | A2A9-6 | BC6 | AACAAGAC |
| CD43 | MEM-59 | BC70 | GTAGGACC |
| CD44 | F10-44-2 | BC2 | TATCCCTT |
| CD45 | HI30 | BC50 | ATAGGGGA |
| CD45RA | HI100 | BC39 | GGAGCCAA |
| CD45RB | MEM55 | BC23 | GCATTCTG |
| CD47 | B6H12-2 | BC85 | GATAGGGT |
| CD48 | MEM-102 | BC69 | GACCACTC |
| CD49D | 9F10 | BC107 | ACCTTGTG |
| CD5 | UCHT2 | BC87 | CAGATCCG |
| CD50 | MEM-171 | BC76 | ACTCTCAC |
| CD52 | HI186 | BC15 | GACTGGGA |
| CD53 | MEM-53 | BC16 | TGCATGTC |
| CD54 | 1H4 | BC9 | GAAAGTCA |
| CD55 | F4-29D9 | BC88 | CAGTCAGT |
| CD59 | MEM-43 | BC29 | GAGGTTAG |
| CD62P | AK4 | BC74 | ATGACTGC |

| | | | |
|---|---|---|---|
| CD64 | 10.1 | BC105 | GCTGAACC |
| CD69 | FN50 | BC47 | AAGCATAG |
| CD7 | 4H9 | BC83 | GATTGTGC |
| CD71 | CY1G4 | BC67 | GCACTTAG |
| CD72 | 3F3 | BC86 | GGTTTACC |
| CD8 | SK1 | BC27 | CTCAGATG |
| CD82 | C33 | BC22 | AACCTTCC |
| CD84 | CD84-1-21 | BC56 | AGTTATCG |
| CD86 | IT2-2 | BC33 | TCTGCTCC |
| CD9 | MEM-61 | BC46 | ACCAGTAC |
| HLA-ABC | W6-32 | BC8 | ATTGGCAC |
| HLA-DR | L243 | BC35 | CCAGCATG |
| SIGLEC | S7-7 | BC82 | AAGTAGCT |
| TCRb | MEM-262 | BC30 | GCGCAACT |
| ACTB | 137CT26-1-1 | BC68 | TCGTAACG |
| mIgG1 | MOPC-21 | BC45 | TTTGGAAG |
| mIgG2a | MOPC-173 | BC31 | CTACTCGC |
| mIgG2b | MPC-11 | BC44 | GTATCGGT |

### **Supplementary Table 2**. List of oligo sequences

| name | 5' mod | 3' mod | sequence |
|---|---|---|---|
| PixA_PL | Phosphate | | TGGTTAGGCGAATTGGACGAGGNNNNNNNNNNNNNNNNNNNNNNNNNGGAACTCGAGGAATGTAAGACC |
| PixA_PLT | | | CCTCGTCCAATTCGCCTAACCAGGTCTTACATTCCTCGAGTTCC |
| PixB_PL | Phosphate | | TCTTTCCCTACACGACGCTCTTCCGATCTNNNNNNNNNNNNNNNNNNNNNNNNNCTGCCTGTGTATCCTGATGTTG |
| PixB_PLT | | | AAGAGCGTCGTGTAGGGAAAGACAACATCAGGATACACAGGCAG |
| PixA_RCaprimer | | | CTCGTCCAATTCGCCTAACCAG |
| PixB_RCAprimer | | | AGAGCGTCGTGTAGGGAAAGAC |
| A_GFP | Phosphate | | CTGCCTGTGTATCCTGATGTTGGTTAGGCGAATTGGACGAGG |
| B_GFP | | | mUmUmUmUmUTCTTTCCCTACACGACGCTCTTCCG |
| ILM_p5_PCR | | | AATGATACGGCGACCACCGAGATCTACAC[Index_8nt]ACACTCTTTCCCTACACGACG*C*T*C |
| ILM_p7_PCR | | | CAAGCAGAAGACGGCATACGAGAT[Index_8nt]GTGACTGGAGTTCAGAC*G*T*G |
| BC1 | Phosphate | Azido-PEG4 | GGAACTCGAGGAATGTAAGACCNNNNNNNNNNNNTCCCTTGCAGATCGGAAGAGCACACGTCTGAACTCCAGTCAC |
| BC2 | Phosphate | Azido-PEG4 | GGAACTCGAGGAATGTAAGACCNNNNNNNNNNNNTATCCCTTAGATCGGAAGAGCACACGTCTGAACTCCAGTCAC |
| BC4 | Phosphate | Azido-PEG4 | GGAACTCGAGGAATGTAAGACCNNNNNNNNNNNNGCCGGACGAGATCGGAAGAGCACACGTCTGAACTCCAGTCAC |
| BC5 | Phosphate | Azido-PEG4 | GGAACTCGAGGAATGTAAGACCNNNNNNNNNNNNTTCTGGGTAGATCGGAAGAGCACACGTCTGAACTCCAGTCAC |
| BC6 | Phosphate | Azido-PEG4 | GGAACTCGAGGAATGTAAGACCNNNNNNNNNNNNAACAAGACAGATCGGAAGAGCACACGTCTGAACTCCAGTCAC |
| BC8 | Phosphate | Azido-PEG4 | GGAACTCGAGGAATGTAAGACCNNNNNNNNNNNNATTGGCACAGATCGGAAGAGCACACGTCTGAACTCCAGTCAC |
| BC9 | Phosphate | Azido-PEG4 | GGAACTCGAGGAATGTAAGACCNNNNNNNNNNNNGAAAGTCAAGATCGGAAGAGCACACGTCTGAACTCCAGTCAC |
| BC11 | Phosphate | Azido-PEG4 | GGAACTCGAGGAATGTAAGACCNNNNNNNNNNNNGTCTTTGTAGATCGGAAGAGCACACGTCTGAACTCCAGTCAC |
| BC12 | Phosphate | Azido-PEG4 | GGAACTCGAGGAATGTAAGACCNNNNNNNNNNNNGTTGTCCGAGATCGGAAGAGCACACGTCTGAACTCCAGTCAC |
| BC13 | Phosphate | Azido-PEG4 | GGAACTCGAGGAATGTAAGACCNNNNNNNNNNNNTCAACGCTAGATCGGAAGAGCACACGTCTGAACTCCAGTCAC |
| BC14 | Phosphate | Azido-PEG4 | GGAACTCGAGGAATGTAAGACCNNNNNNNNNNNNGTGCATTCAGATCGGAAGAGCACACGTCTGAACTCCAGTCAC |
| BC15 | Phosphate | Azido-PEG4 | GGAACTCGAGGAATGTAAGACCNNNNNNNNNNNNGACTGGGAAGATCGGAAGAGCACACGTCTGAACTCCAGTCAC |
| BC16 | Phosphate | Azido-PEG4 | GGAACTCGAGGAATGTAAGACCNNNNNNNNNNNNTGCATGTCAGATCGGAAGAGCACACGTCTGAACTCCAGTCAC |
| BC17 | Phosphate | Azido-PEG4 | GGAACTCGAGGAATGTAAGACCNNNNNNNNNNNNAAGTCGTGAGATCGGAAGAGCACACGTCTGAACTCCAGTCAC |
| BC18 | Phosphate | Azido-PEG4 | GGAACTCGAGGAATGTAAGACCNNNNNNNNNNNNACATTGACAGATCGGAAGAGCACACGTCTGAACTCCAGTCAC |
| BC19 | Phosphate | Azido-PEG4 | GGAACTCGAGGAATGTAAGACCNNNNNNNNNNNNTGATAGAAAGATCGGAAGAGCACACGTCTGAACTCCAGTCAC |
| BC20 | Phosphate | Azido-PEG4 | GGAACTCGAGGAATGTAAGACCNNNNNNNNNNNNATTCGCCTAGATCGGAAGAGCACACGTCTGAACTCCAGTCAC |
| BC22 | Phosphate | Azido-PEG4 | GGAACTCGAGGAATGTAAGACCNNNNNNNNNNNNAACCTTCCAGATCGGAAGAGCACACGTCTGAACTCCAGTCAC |
| BC23 | Phosphate | Azido-PEG4 | GGAACTCGAGGAATGTAAGACCNNNNNNNNNNNNGCATTCTGAGATCGGAAGAGCACACGTCTGAACTCCAGTCAC |
| BC25 | Phosphate | Azido-PEG4 | GGAACTCGAGGAATGTAAGACCNNNNNNNNNNNNTCAGGGTGAGATCGGAAGAGCACACGTCTGAACTCCAGTCAC |
| BC26 | Phosphate | Azido-PEG4 | GGAACTCGAGGAATGTAAGACCNNNNNNNNNNNNCTACGACGAGATCGGAAGAGCACACGTCTGAACTCCAGTCAC |
| BC27 | Phosphate | Azido-PEG4 | GGAACTCGAGGAATGTAAGACCNNNNNNNNNNNNCTCAGATGAGATCGGAAGAGCACACGTCTGAACTCCAGTCAC |
| BC29 | Phosphate | Azido-PEG4 | GGAACTCGAGGAATGTAAGACCNNNNNNNNNNNNGAGGTTAGAGATCGGAAGAGCACACGTCTGAACTCCAGTCAC |
| BC30 | Phosphate | Azido-PEG4 | GGAACTCGAGGAATGTAAGACCNNNNNNNNNNNNGCGCAACTAGATCGGAAGAGCACACGTCTGAACTCCAGTCAC |
| BC31 | Phosphate | Azido-PEG4 | GGAACTCGAGGAATGTAAGACCNNNNNNNNNNNNCTACTCGCAGATCGGAAGAGCACACGTCTGAACTCCAGTCAC |
| BC32 | Phosphate | Azido-PEG4 | GGAACTCGAGGAATGTAAGACCNNNNNNNNNNNNACTGTGTCAGATCGGAAGAGCACACGTCTGAACTCCAGTCAC |
| BC33 | Phosphate | Azido-PEG4 | GGAACTCGAGGAATGTAAGACCNNNNNNNNNNNNTCTGCTCCAGATCGGAAGAGCACACGTCTGAACTCCAGTCAC |
| BC34 | Phosphate | Azido-PEG4 | GGAACTCGAGGAATGTAAGACCNNNNNNNNNNNNAGGATGTTAGATCGGAAGAGCACACGTCTGAACTCCAGTCAC |
| BC35 | Phosphate | Azido-PEG4 | GGAACTCGAGGAATGTAAGACCNNNNNNNNNNNNCCAGCATGAGATCGGAAGAGCACACGTCTGAACTCCAGTCAC |
| BC36 | Phosphate | Azido-PEG4 | GGAACTCGAGGAATGTAAGACCNNNNNNNNNNNNAGCTACTAAGATCGGAAGAGCACACGTCTGAACTCCAGTCAC |
| BC37 | Phosphate | Azido-PEG4 | GGAACTCGAGGAATGTAAGACCNNNNNNNNNNNNCCGATATCAGATCGGAAGAGCACACGTCTGAACTCCAGTCAC |
| BC38 | Phosphate | Azido-PEG4 | GGAACTCGAGGAATGTAAGACCNNNNNNNNNNNNGAATCCCGAGATCGGAAGAGCACACGTCTGAACTCCAGTCAC |
| BC39 | Phosphate | Azido-PEG4 | GGAACTCGAGGAATGTAAGACCNNNNNNNNNNNNGGAGCCAAAGATCGGAAGAGCACACGTCTGAACTCCAGTCAC |
| BC40 | Phosphate | Azido-PEG4 | GGAACTCGAGGAATGTAAGACCNNNNNNNNNNNNAGAGGCTCAGATCGGAAGAGCACACGTCTGAACTCCAGTCAC |
| BC42 | Phosphate | Azido-PEG4 | GGAACTCGAGGAATGTAAGACCNNNNNNNNNNNNCTGACCATAGATCGGAAGAGCACACGTCTGAACTCCAGTCAC |
| BC44 | Phosphate | Azido-PEG4 | GGAACTCGAGGAATGTAAGACCNNNNNNNNNNNNGTATCGGTAGATCGGAAGAGCACACGTCTGAACTCCAGTCAC |
| BC45 | Phosphate | Azido-PEG4 | GGAACTCGAGGAATGTAAGACCNNNNNNNNNNNNTTTGGAAGAGATCGGAAGAGCACACGTCTGAACTCCAGTCAC |
| BC46 | Phosphate | Azido-PEG4 | GGAACTCGAGGAATGTAAGACCNNNNNNNNNNNNACCAGTACAGATCGGAAGAGCACACGTCTGAACTCCAGTCAC |
| BC47 | Phosphate | Azido-PEG4 | GGAACTCGAGGAATGTAAGACCNNNNNNNNNNNNAAGCATAGAGATCGGAAGAGCACACGTCTGAACTCCAGTCAC |
| BC48 | Phosphate | Azido-PEG4 | GGAACTCGAGGAATGTAAGACCNNNNNNNNNNNNCTGTAGGAAGATCGGAAGAGCACACGTCTGAACTCCAGTCAC |
| BC49 | Phosphate | Azido-PEG4 | GGAACTCGAGGAATGTAAGACCNNNNNNNNNNNNATTTCGAGAGATCGGAAGAGCACACGTCTGAACTCCAGTCAC |
| BC50 | Phosphate | Azido-PEG4 | GGAACTCGAGGAATGTAAGACCNNNNNNNNNNNNATAGGGGAAGATCGGAAGAGCACACGTCTGAACTCCAGTCAC |
| BC51 | Phosphate | Azido-PEG4 | GGAACTCGAGGAATGTAAGACCNNNNNNNNNNNNAACGGCTAAGATCGGAAGAGCACACGTCTGAACTCCAGTCAC |
| BC53 | Phosphate | Azido-PEG4 | GGAACTCGAGGAATGTAAGACCNNNNNNNNNNNNTCCCAGTGAGATCGGAAGAGCACACGTCTGAACTCCAGTCAC |
| BC54 | Phosphate | Azido-PEG4 | GGAACTCGAGGAATGTAAGACCNNNNNNNNNNNNTACTCTTGAGATCGGAAGAGCACACGTCTGAACTCCAGTCAC |
| BC56 | Phosphate | Azido-PEG4 | GGAACTCGAGGAATGTAAGACCNNNNNNNNNNNNAGTTATCGAGATCGGAAGAGCACACGTCTGAACTCCAGTCAC |
| BC57 | Phosphate | Azido-PEG4 | GGAACTCGAGGAATGTAAGACCNNNNNNNNNNNNGTGAGTAGAGATCGGAAGAGCACACGTCTGAACTCCAGTCAC |
| BC58 | Phosphate | Azido-PEG4 | GGAACTCGAGGAATGTAAGACCNNNNNNNNNNNNCATGGGCAAGATCGGAAGAGCACACGTCTGAACTCCAGTCAC |
| BC59 | Phosphate | Azido-PEG4 | GGAACTCGAGGAATGTAAGACCNNNNNNNNNNNNAGGGCAGTAGATCGGAAGAGCACACGTCTGAACTCCAGTCAC |
| BC61 | Phosphate | Azido-PEG4 | GGAACTCGAGGAATGTAAGACCNNNNNNNNNNNNCCTAAGAAAGATCGGAAGAGCACACGTCTGAACTCCAGTCAC |
| BC62 | Phosphate | Azido-PEG4 | GGAACTCGAGGAATGTAAGACCNNNNNNNNNNNNGTTCAGACAGATCGGAAGAGCACACGTCTGAACTCCAGTCAC |
| BC63 | Phosphate | Azido-PEG4 | GGAACTCGAGGAATGTAAGACCNNNNNNNNNNNNAGCCCGAAAGATCGGAAGAGCACACGTCTGAACTCCAGTCAC |
| BC64 | Phosphate | Azido-PEG4 | GGAACTCGAGGAATGTAAGACCNNNNNNNNNNNNTGGGGCTTAGATCGGAAGAGCACACGTCTGAACTCCAGTCAC |
| BC65 | Phosphate | Azido-PEG4 | GGAACTCGAGGAATGTAAGACCNNNNNNNNNNNNCACACGGTAGATCGGAAGAGCACACGTCTGAACTCCAGTCAC |
| BC67 | Phosphate | Azido-PEG4 | GGAACTCGAGGAATGTAAGACCNNNNNNNNNNNNGCACTTAGAGATCGGAAGAGCACACGTCTGAACTCCAGTCAC |
| BC68 | Phosphate | Azido-PEG4 | GGAACTCGAGGAATGTAAGACCNNNNNNNNNNNNTCGTAACGAGATCGGAAGAGCACACGTCTGAACTCCAGTCAC |

| | | | |
|---|---|---|---|
| BC69 | Phosphate | Azido-PEG4 | GGAACTCGAGGAATGTAAGACCNNNNNNNNNNGACCACTCAGATCGGAAGAGCACACGTCTGAACTCCAGTCAC |
| BC70 | Phosphate | Azido-PEG4 | GGAACTCGAGGAATGTAAGACCNNNNNNNNNNGTAGGACCAGATCGGAAGAGCACACGTCTGAACTCCAGTCAC |
| BC72 | Phosphate | Azido-PEG4 | GGAACTCGAGGAATGTAAGACCNNNNNNNNNNCTTGCACCAGATCGGAAGAGCACACGTCTGAACTCCAGTCAC |
| BC73 | Phosphate | Azido-PEG4 | GGAACTCGAGGAATGTAAGACCNNNNNNNNNNCTCAAGAGAGATCGGAAGAGCACACGTCTGAACTCCAGTCAC |
| BC74 | Phosphate | Azido-PEG4 | GGAACTCGAGGAATGTAAGACCNNNNNNNNNNATGACTGCAGATCGGAAGAGCACACGTCTGAACTCCAGTCAC |
| BC76 | Phosphate | Azido-PEG4 | GGAACTCGAGGAATGTAAGACCNNNNNNNNNNACTCTCACAGATCGGAAGAGCACACGTCTGAACTCCAGTCAC |
| BC77 | Phosphate | Azido-PEG4 | GGAACTCGAGGAATGTAAGACCNNNNNNNNNNTCCGTAACAGATCGGAAGAGCACACGTCTGAACTCCAGTCAC |
| BC79 | Phosphate | Azido-PEG4 | GGAACTCGAGGAATGTAAGACCNNNNNNNNNNTTGTCCAAAGATCGGAAGAGCACACGTCTGAACTCCAGTCAC |
| BC80 | Phosphate | Azido-PEG4 | GGAACTCGAGGAATGTAAGACCNNNNNNNNNNGCTATTGAAGATCGGAAGAGCACACGTCTGAACTCCAGTCAC |
| BC82 | Phosphate | Azido-PEG4 | GGAACTCGAGGAATGTAAGACCNNNNNNNNNNAAGTAGCTAGATCGGAAGAGCACACGTCTGAACTCCAGTCAC |
| BC83 | Phosphate | Azido-PEG4 | GGAACTCGAGGAATGTAAGACCNNNNNNNNNNGATTGTGCAGATCGGAAGAGCACACGTCTGAACTCCAGTCAC |
| BC84 | Phosphate | Azido-PEG4 | GGAACTCGAGGAATGTAAGACCNNNNNNNNNNTTTCTGGTAGATCGGAAGAGCACACGTCTGAACTCCAGTCAC |
| BC85 | Phosphate | Azido-PEG4 | GGAACTCGAGGAATGTAAGACCNNNNNNNNNNGATAGGGTAGATCGGAAGAGCACACGTCTGAACTCCAGTCAC |
| BC86 | Phosphate | Azido-PEG4 | GGAACTCGAGGAATGTAAGACCNNNNNNNNNNGGTTTACCAGATCGGAAGAGCACACGTCTGAACTCCAGTCAC |
| BC87 | Phosphate | Azido-PEG4 | GGAACTCGAGGAATGTAAGACCNNNNNNNNNNCAGATCCGAGATCGGAAGAGCACACGTCTGAACTCCAGTCAC |
| BC88 | Phosphate | Azido-PEG4 | GGAACTCGAGGAATGTAAGACCNNNNNNNNNNCAGTCAGTAGATCGGAAGAGCACACGTCTGAACTCCAGTCAC |
| BC101 | Phosphate | Azido-PEG4 | GGAACTCGAGGAATGTAAGACCNNNNNNNNNNAAAGCAACAGATCGGAAGAGCACACGTCTGAACTCCAGTCAC |
| BC102 | Phosphate | Azido-PEG4 | GGAACTCGAGGAATGTAAGACCNNNNNNNNNNCAATCGGCAGATCGGAAGAGCACACGTCTGAACTCCAGTCAC |
| BC104 | Phosphate | Azido-PEG4 | GGAACTCGAGGAATGTAAGACCNNNNNNNNNNCACGTTTCAGATCGGAAGAGCACACGTCTGAACTCCAGTCAC |
| BC105 | Phosphate | Azido-PEG4 | GGAACTCGAGGAATGTAAGACCNNNNNNNNNNGCTGAACCAGATCGGAAGAGCACACGTCTGAACTCCAGTCAC |
| BC107 | Phosphate | Azido-PEG4 | GGAACTCGAGGAATGTAAGACCNNNNNNNNNNACCTTGTGAGATCGGAAGAGCACACGTCTGAACTCCAGTCAC |
| BC108 | Phosphate | Azido-PEG4 | GGAACTCGAGGAATGTAAGACCNNNNNNNNNNACTCGGAAAGATCGGAAGAGCACACGTCTGAACTCCAGTCAC |
| BC109 | Phosphate | Azido-PEG4 | GGAACTCGAGGAATGTAAGACCNNNNNNNNNNCTTCTTGAAGATCGGAAGAGCACACGTCTGAACTCCAGTCAC |
| BC110 | Phosphate | Azido-PEG4 | GGAACTCGAGGAATGTAAGACCNNNNNNNNNNCCAGACACAGATCGGAAGAGCACACGTCTGAACTCCAGTCAC |
